# Early detection of cerebrovascular pathology and protective antiviral immunity by MRI

**DOI:** 10.1101/2021.10.21.465276

**Authors:** Li Liu, Stephen Dodd, Ryan D. Hunt, Nikorn Pothayee, Tatjana Atanasijevic, Nadia Bouraoud, Dragan Maric, E. Ashley Moseman, Selamawit Gossa, Dorian B. McGavern, Alan P. Koretsky

## Abstract

Central nervous system (CNS) infections are a major cause of human morbidity and mortality worldwide. Even patients that survive CNS infections can have lasting neurological dysfunction resulting from immune and pathogen induced pathology. Developing approaches to noninvasively track pathology and immunity in the infected CNS is crucial for patient management and development of new therapeutics. Here, we develop novel MRI-based approaches to monitor virus-specific CD8+ T cells and their relationship to cerebrovascular pathology in the living brain. We studied a relevant murine model in which a neurotropic virus (vesicular stomatitis virus) was introduced intranasally and then entered the brain via olfactory sensory neurons – a route exploited by many pathogens in humans. Using T2*-weighted high-resolution MRI, we identified small cerebral microbleeds as the earliest form of pathology associated with viral entry into the brain. Mechanistically, these microbleeds occurred in the absence of peripheral immune cells and were associated with infection of vascular endothelial cells. We monitored the adaptive response to this infection by developing methods to iron label and track individual virus specific CD8+ T cells by MRI. Transferred antiviral T cells were detected in the brain within a day of infection and were able to reduce cerebral microbleeds. These data demonstrate the utility of MRI in detecting the earliest pathological events in the virally infected CNS as well as the therapeutic potential of antiviral T cells in mitigating this pathology.

## Introduction

The central nervous system (CNS) is protected by several physical barriers, however, a variety of neurotropic viruses from different families are able to infect the CNS (Swanson & McGavern, 2015). Viruses use multiple strategies to access the CNS. Some can infect the brain through the blood brain barrier (BBB) directly, whereas others enter the peripheral nervous system and use axonal transport along motor and olfactory neurons. Infection of hematopoietic cells is also used to enter the CNS (McGavern & Kang, 2011).

One common pathology associated with viral infection is bleeding. Two major causes of virus-induced CNS bleeding are direct killing of infected cells and immune-mediated damage (Ludlow et al., 2016). Neural and glial inflammatory reactions play an important role in the defense against a viral infection (Chhatbar et al., 2018), as do innate and adaptive immune cells recruited from the periphery (Manglani & McGavern, 2017). Antiviral T cells recruited into the brain in response to infection can be both helpful and harmful (Klein & Hunter, 2017). For example, studies have shown that antiviral memory T cells adoptively transferred into mice persistently infected from birth with lymphocytic choriomeningitis virus (LCMV) can clear virus from the brain without causing immunopathology (Herz et al., 2015). However, pathogen-specific CD8 T cells can also trigger fatal vascular breakdown, which has been observed during cerebral malaria (Belnoue et al., 2002; Riggle et al., 2020; Swanson et al., 2016), viral meningitis (Kim et al., 2009), and SARS-CoV-2 infection (Schwabenland et al., 2021). Thus, immunotherapies designed to treat CNS pathogens face more challenges than those focused on peripheral infections due to the sensitivity of the CNS compartment as well as the unique features and functions of cerebrovasculature.

Magnetic resonance imaging (MRI) has been very useful for the detection of pathological changes in the CNS during viral infection (Lee et al., 2020; Poyiadji et al., 2020). Microbleeds can be detected sensitively as punctate hypointensity spots on T2*-weighted MRI (Griffin et al., 2019). In post-mortem brain tissue from COVID patients, microbleeds were detected by T2*-weighted MRI, which guided the histology studies of cerebrovascular pathology and associated inflammation (Lee et al., 2020). Recently, there has been growing interest in using MRI to track cell movements, even at the single cell level (Pothayee et al., 2017). Noninvasive MRI based tracking of antiviral T cells in relation to microbleeds should provide additional insights into the mechanisms that result in CNS vascular bleeding during different types of infections.

In this study, we used intranasal infection of mice with vesicular stomatitis virus (VSV) as a model (Sabin & Olitsky, 1937) to explore cerebrovascular pathology and antiviral T cells by MRI. The olfactory bulb (OB) serves as the CNS entry point for many neurotropic viruses inhaled into the nose. For example, VSV can infect olfactory sensory neurons that project fibers from the nasal airway to the OB, enabling viral transit within axon fibers and CNS entry (Chhatbar et al., 2018; Moseman et al., 2020). The main goal of this study was to noninvasively define the temporal and spatial relationship between cerebrovascular breakdown and CD8 T cell infiltration in the virus-infected brain. More specifically, our aims were to: (i) characterize early stage microbleeds by T2*-weighted MRI as an neuroimaging marker for CNS viral infection; (ii) determine whether VSV-induced cerebrovascular damage was caused by the virus or infiltrating immune cells; (iii) evaluate whether adoptively transferred virus-specific CD8 T cells could prevent cerebrovascular damage; (iv) develop an MRI technique to label and track antiviral CD8 T cells; (v) define the relationship between microbleeds and CD8 T cells by MRI. Our results demonstrate that VSV promotes extensive bleeding in the OB and other brain regions. Bleeding occurred even in the absence of peripheral immune infiltration, suggesting that the virus alone (or, in combination with brain resident cells) can cause vascular breakdown. Spatially, virus-specific CD8 T cells were observed in brain regions with and without bleeding, and vascular pathology was reduced in mice supplemented with these antiviral cells.

## Results

### Detection of microbleeds in the VSV-infected turbinates and brain by MRI

Upon intranasal VSV inoculation, immune cell accumulation in the turbinates and olfactory bulbs (OB) has been reported (Chhatbar et al., 2018; Moseman et al., 2020). To determine how VSV infection affects vessel integrity, high resolution T2* weighted MRI was used to detect bleeding at different time points post-infection (Fig. 1). Large areas of bleeding were readily detected in the turbinates at day 4 (Fig 1A; S2A). The amount of bleeding, as indicated by hypointense areas (red arrows), increased from day 4 to 11. Two major sites of bleeding were also detected by MRI in the OB (Fig 1B-D; S2B). Bleeding was observed at the edge of the bulb, in the olfactory nerve layer (ONL) and glomerular layer (GL), and at the center of the bulb near the granule cell layer (GCL). The *in-vivo* and *ex-vivo* images on each day were obtained from the same mouse before and after perfusion. In the *in-vivo* images (upper panel), the asymmetric hypointense region at the edge of the OB indicates the bleeding. As early as day 4, *in-vivo* MRI revealed microbleeds and vessel rupture starting in the ONL and GL of the OB (red arrows; Fig 1B, C; S2B). *Ex-vivo* high-resolution MRI detected punctate hypointensities in the ONL and GL on day 4, which confirmed the results of *in-vivo* MRI. On day 6, punctate hypointensities near the center of the OB were also detected by *in-vivo* MRI (Fig 1D). The number of microbleeds in the OB increased from day 4 to 11. The increasing amount of hypointensities in the turbinates and OB were quantified as a function of time (Fig 1E). OB viral titers peaked on day 6 before dropping to undetectable levels by day 11 (Fig 1F).

**Figure 1.**
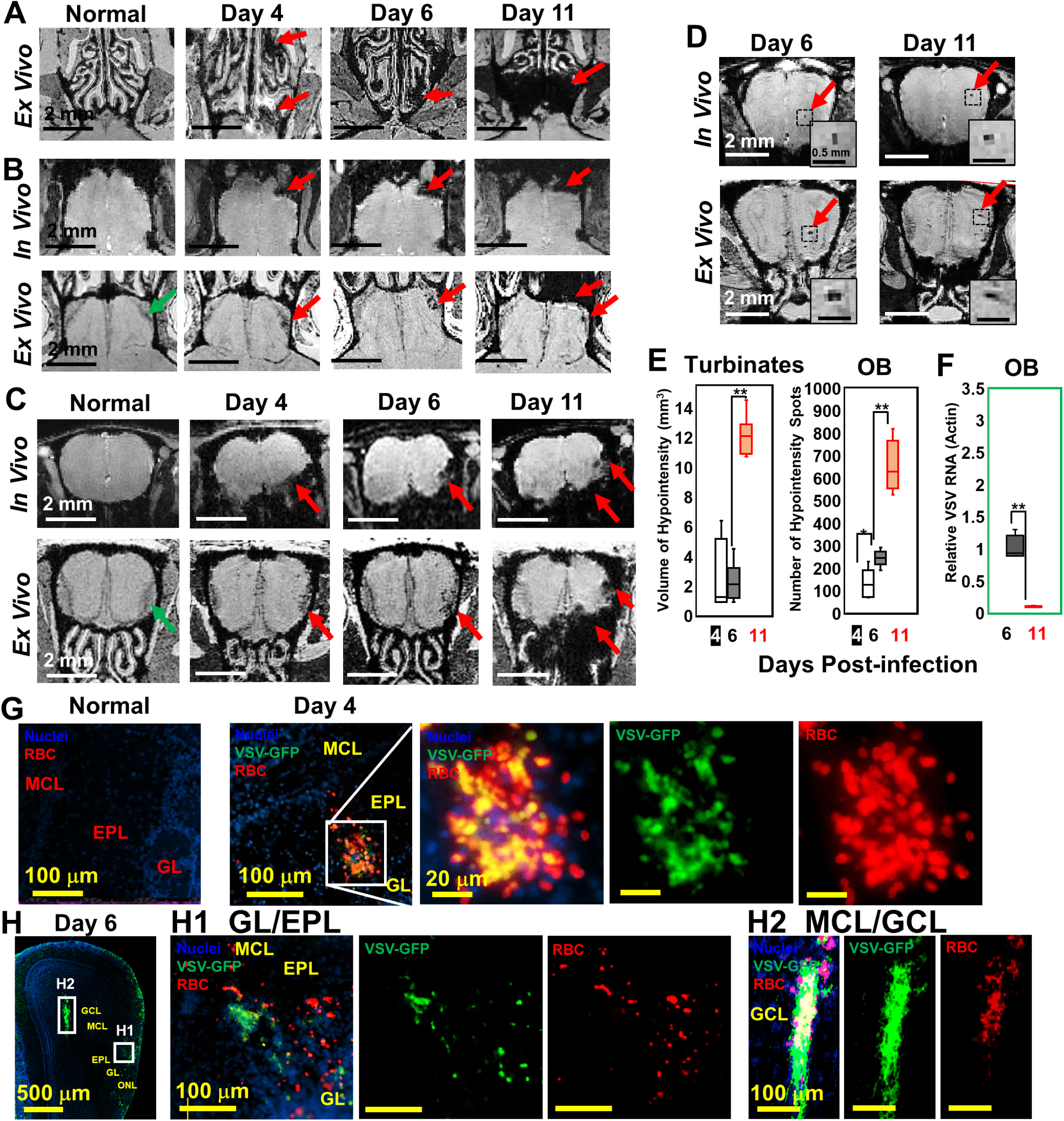
MRI detected microbleeds in turbinates and OB since day 4 post-infection. (A) *Ex-vivo* MRI of turbinates (axial view) from a normal mouse and VSV-infected mice on day 4, 6, and 11 post-infection. (B) Axial view and (C) coronal view of *in-vivo* (upper panel) and *ex-vivo* (lower panel) MR images of OB. The coronal view was sectioned from the bleeding site on the axial view, as indicated by red arrows. (D) At the center of the OB, microbleeds were detected since around day 6 by *in-vivo* MRI. The inserted figures were the enlarged views of the framed bleeding sites in the full views. The scale bar in the full view: 2 mm. The scale bar in the inserted view: 0.5 mm (D). (E) Quantification of volume of hypointensity in the turbinates and numbers of hypointensity spots in the OB on day 4, 6, and 11 (N = 6). *, p = 0.046; **, p < 0.0001. (F) Viral titers, represented as relative to actin RNA, at the OB on day 6 and 11 (N = 6). **, p < 0.0001. (G, H) IHC study to show microbleeds and VSV in the OB of VSV-infected mice on day 4 (G) and 6 (H). EPL, external plexiform layer; GCL, granule cell layer; GL, glomerular layer; MCL, mitral cell layer; ONL, olfactory nerve layer. Blue, DAPI for nuclei; Green, VSV-GFP; Red, Alexa Fluor 647-conjugated anti-Ter119 for RBCs.

Immunohistochemical (IHC) analyses verified the MRI results by showing that the ONL, GL and GCL were the major sites of bleeding (Fig 1G, H; S2D, E). These are also the sites where VSV entered the brain and replicated. On day 4, IHC showed microbleeds and VSV in the ONL, GL and external plexiform layer (EPL) of the OB (Fig 1G, Fig S2D). On day 6, microbleeds and VSV were shown at the mitral cell layer (MCL) (Fig 1 H1; S2E) and center of the OB (Fig 1 H2). Most of the virus was localized in ONL, GL and the center of the OB on day when viral titer was the highest (Fig 1H). Thus, microbleeds largely co-localized with sites of VSV replication in the brain.

Although VSV replication is usually contained within the OB (Detje et al., 2009), we detected vessel breakdown and microbleeds in other parts of the brain by MRI (Fig 2A, B; more views shown in Fig S3A, B). Punctate hypointensities were detected from frontal brain to midbrain beginning at day 4. In most of the infected mice, microbleeds were detected in the orbital frontal cortex area and somatomotor areas of the cortex around day 4. On day 6, microbleeds were detected in the caudoputamen (CP) and thalamus (TH). From day 8 to 11, microbleeds were observed in the midbrain (MB). Through IHC study, microbleeds and VSV were observed at the forebrain on day 6 and in the midbrain on day 11 (Fig 2D; S3C, D). The increasing amount of hypointensities in the brain (not including the OB) were quantified as a function of time (Fig 2C). For the OB and other brain regions, sites of bleeding were enumerated without accounting for size. This was done to assess the number of new bleeding regions that developed. Foci of bleeding (defined as spots/mm^3^) was ∼100-fold greater in the OB than the brain. For these calculations, the size of mouse OB was assumed to be ∼2% of the brain by volume (McGann, 2017). The elevated bleeding in the OB is consistent with the fact that VSV is most abundant in this brain region following intranasal inoculation.

**Figure 2.**
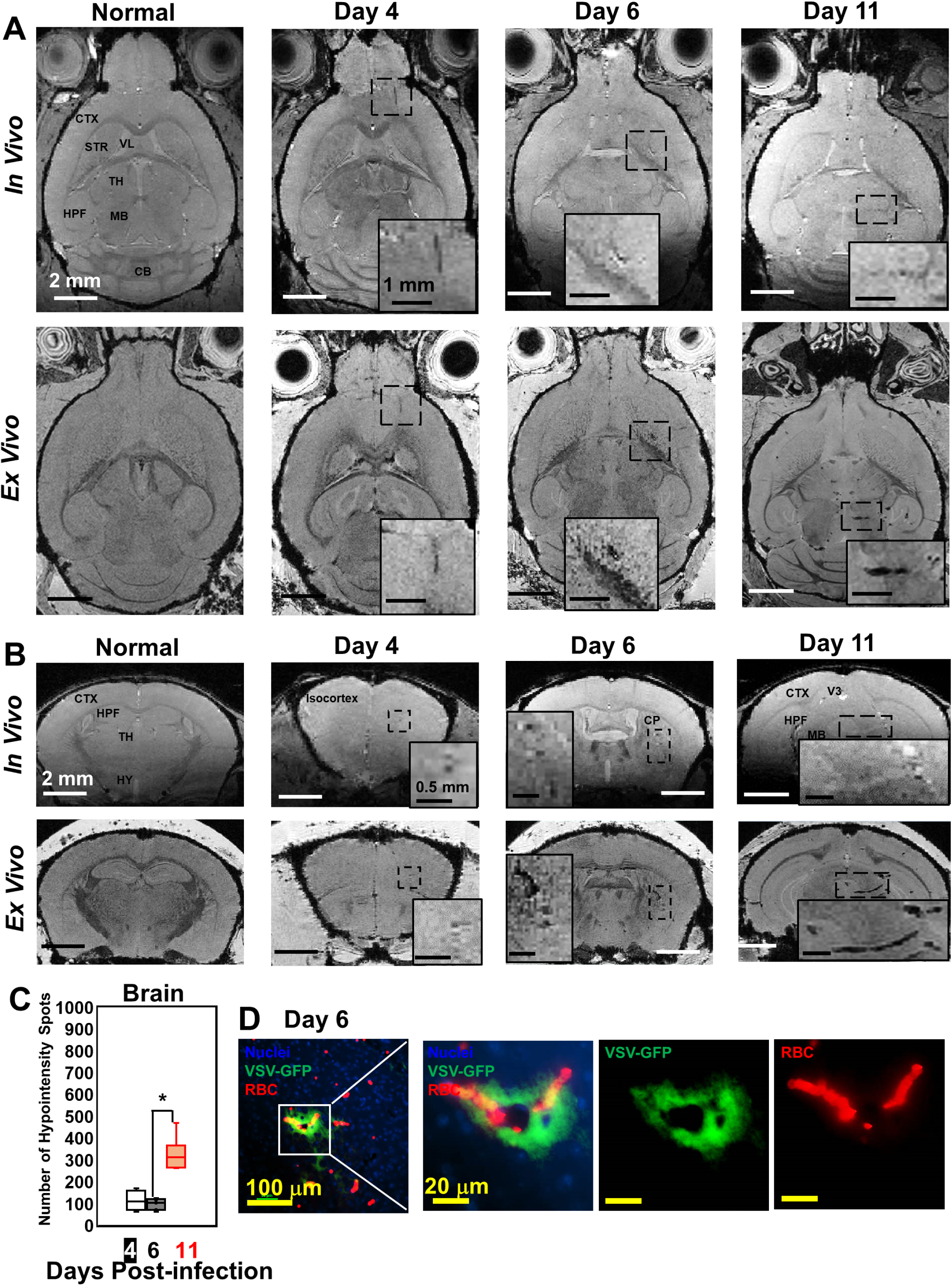
MRI monitored vessel breakdown in the rest of brain. (A) Axial view and (B) coronal view of *in-vivo* (upper panel) and *ex-vivo* (lower panel) MR images of brain, showing the breakdown of vessels from frontal brain to midbrain. The scale bar in the full view: 2 mm. The scale bar in the inserted view: 1mm (A) and 0.5 mm (B). CB, cerebellum; CP, caudoputamen; CTX, cerebral cortex; HPF, hippocampal formation; HY, hypothalamus; MB, midbrain; STR, striatum; TH, thalamus; V3, third ventricle; VL, lateral ventricle. (C) Quantification of numbers of hypointensity spots in the brain (not including OB) on day 4, 6, and 11 (N = 6). *, p < 0.0001. (D) IHC staining to show microbleeds and VSV in the forebrain on day 6. Blue, DAPI for nuclei; Green, VSV-GFP; Red, Alexa Fluor 647-conjugated anti-Ter119 for RBCs.

### VSV caused vascular endothelium activation and induced cerebral hemorrhage does not require peripheral immune cells

Significantly increased expression of vascular cell adhesion molecule 1 (VCAM-1) and intercellular adhesion molecule 1 (ICAM-1) were found in the OB on day 6 of infection. VCAM-1 and ICAM-1 vessel surface coverage (by using anti-CD31, or anti-PECAM-1, to stain endothelial cells) was around 50% near the ONL/GL/EPL region (Fig 3A) and around 30% near the center of the OB (Fig S4A, C). Upregulation of VCAM-1 and ICAM-1 were also found in many other locations in the midbrain and hindbrain on day 6, e.g., thalamus (Fig S4B, C).

**Figure 3.**
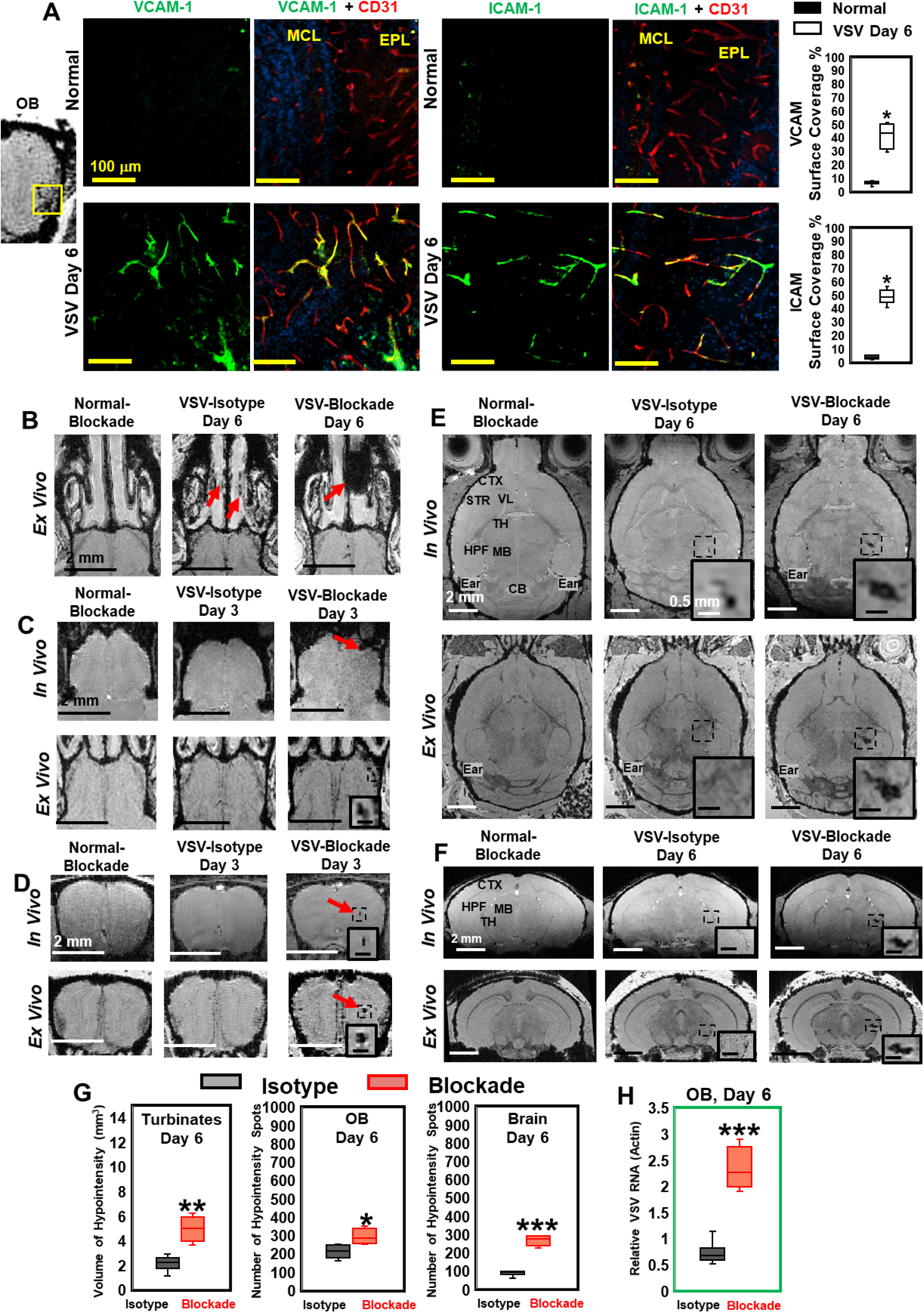
VSV infection caused vascular endothelium activation and blocking immune cell infiltration by anti-LFA-1/VLA-4 antibodies promoted bleeding in the turbinates, OB, and brain. (A) IHC staining showed increased expression of endothelial VCAM-1 and ICAM-1 in the OB on day 6 of infection. N = 3 per group. * p < 0.0001. (B) Treatment with anti-LFA-1/VLA-4 blockade antibodies increased bleeding in the turbinates. *Ex-vivo* MRI of the turbinates (axial view) from normal mouse treated with the blockade antibodies, VSV infected mouse treated with isotype control (rat IgG2a) antibody on day 6, and VSV infected mouse treated with the blockade antibodies on day 6. In the GL (C) and the GCL of OB (D), MRI detected bleeding earlier upon treatment with the blockade antibodies. (E) Axial and (F) coronal view of MR images of the brain showed a large area of hemorrhage at the thalamus of anti-LFA-1/VLA-4 treated mouse on day 6. The coronal view was sectioned from the hemorrhage site on the axial view, as shown in the frame. In (E) and (F), scale bar in the full view: 2 mm; in the inserted view: 0.5 mm. (G) Quantification of the volume of bleeding in the turbinates and the numbers of hypointensity spots in the OB and brain on day 6, in the isotype control group and anti-LFA-1/VLA-4 antibodies treated group (N = 4). *, p = 0.039; **, p = 0.005; ***, p < 0.0001. (H) Blocking immune cell infiltration increased viral titers in the OB on day 6 (N = 4). ***, p < 0.0001.

To determine whether the vessel breakdown and microbleeds associated with VSV infection required the peripheral immune system, anti-LFA-1/VLA-4 antibodies were used to block infiltration by peripheral immune cells. LFA-1 and VLA-4 are the ligands for ICAM-1 and VCAM-1, which are expressed on activated T cells, monocytes, and neutrophils (Yusuf-Makagiansar et al., 2002). Upon treatment with anti-LFA-1/VLA-4 blocking antibodies, a greater volume of bleeding was detected in the turbinates on day 6, relative to the two control groups, i.e., normal mice treated with anti-LFA-1/VLA-4 antibodies and VSV infected mice treated with isotype control (rat IgG2a) antibodies (Fig 3B). Within the OB, *in-vivo* MRI revealed bleeding as early as day 3 (Fig 3C, *in-vivo* panel). The earliest time point that *in-vivo* MRI could detect bleeding in the OB of mice treated with isotype control antibodies was on day 4, which was the same as untreated VSV-infected mice (Fig 1B). Blocking immune cell infiltration also increased bleeding at the center of the OB on day 3 as detected by *in-vivo* MRI (Fig 3D), and a large area of hemorrhage was detected in the thalamus on day 6 (Fig 3E, F; S4D-F).

As quantified in Fig 3G, the volume of hypointensity in the turbinates increased from 2.1 ± 0.8 mm^3^ to 5.0 ± 1.0 mm^3^ on day 6 in anti-LFA-1/VLA-4 treated mice, and the number of bleeding spots increased in the brain by ∼2-fold. In addition, blockade of immune infiltration elevated viral titers in the OB by ∼2-3 fold on day 6 (Fig 3H). Thus, VSV induces hemorrhage in the brain without involvement of the peripheral immune system. In fact, inhibiting the peripheral immune response increases viral titers and gives rise to more bleeding.

### VSV infects vascular endothelial cells

VSV was previously shown to enter the brain through olfactory sensory neurons via retrograde transneuronal transport and spread in the CNS via both anterograde and retrograde transport (Beier et al., 2013; Cornish et al., 2001). Although VSV is highly cytopathic, most VSV-infected neurons in the brain are cleared noncytolytically (Moseman et al., 2020). While it is generally known that neurons are infected by VSV, we became interested in whether other cell types supported VSV infection. We assessed if cerebrovascular vascular endothelial cells were infected by the virus given the bleeding observed in the brains of VSV-infected mice. As shown in Fig 4A, B, we detected VSV in endothelial cells comprising cerebral blood vessels, which sometimes showed evidence of bleeding (Fig 4C). These data suggest that bleeding might result from infection of cerebrovascular endothelial cells by VSV.

**Figure 4.**
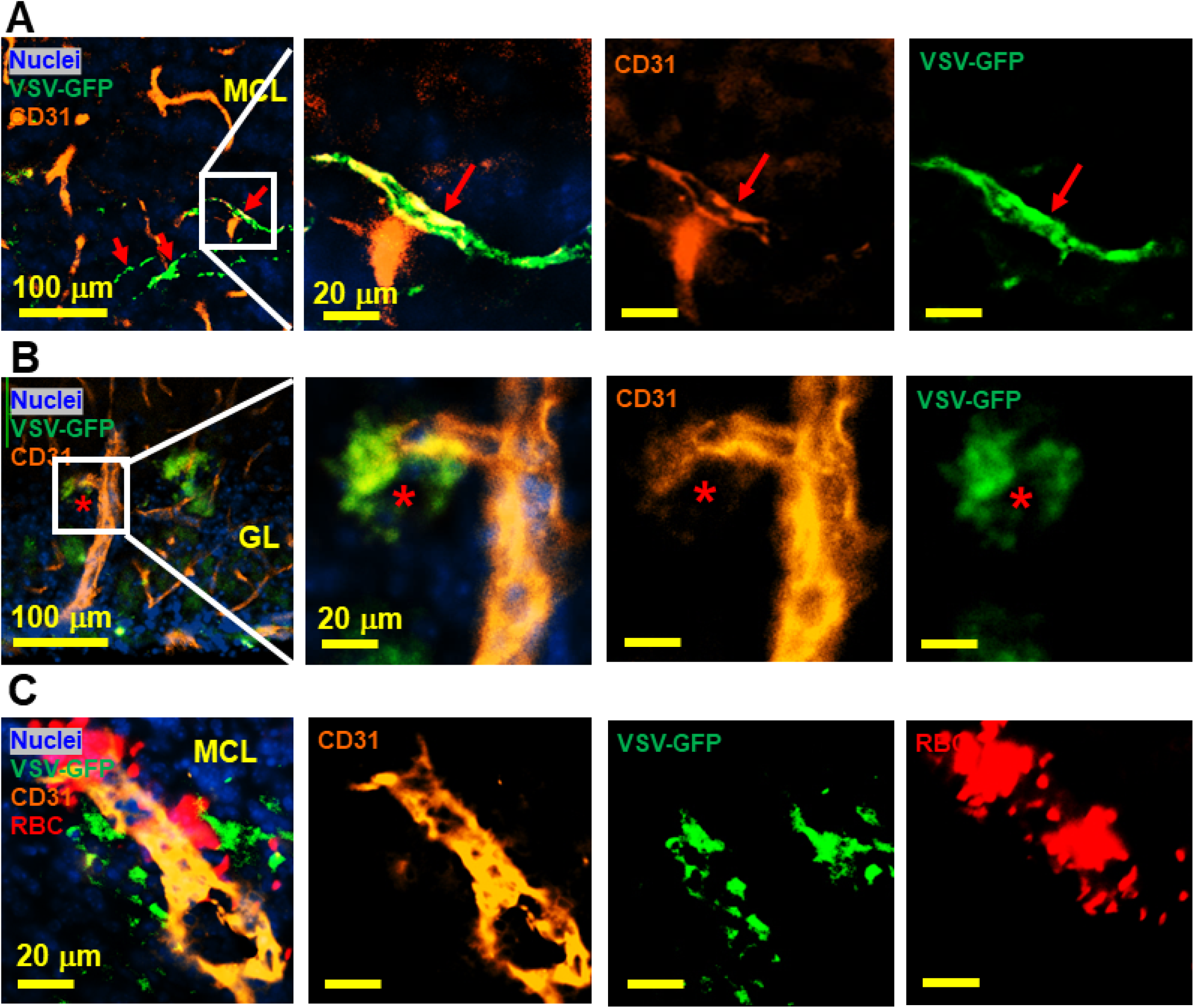
IHC study revealed that VSV can infect vascular endothelial cells. (A, B) Representative views of VSV infected vessels in the MCL and GL, as indicated by the red arrows and red star. Blue, DAPI for nuclei; Green, VSV-GFP; Orange, Alexa Fluor 647 conjugated anti-CD31. (C) Bleedings were observed from VSV infected vessels in the MCL. Orange, Alexa Fluor 594 conjugated anti-CD31; Red, Alexa Fluor 700 conjugated anti-Ter119 for RBCs.

### Adoptive transfer of antiviral CD8 T cells decreases brain bleeding and increases viral clearance

Inhibiting peripheral immune cells led to more brain bleeding. We therefore tested if adoptive transfer of virus-specific CD8 T cells could be used to reduce viral loads and inhibit brain bleeding. 5 x 10^5^ virus-specific mTomato CD8+ OT-I T cells (purity shown in Fig S5) activated overnight with ovalbumin (OVA)_257-264_ peptide were transferred i.v. into VSV-OVA infected mice at two time points. For early-stage transfer, OT-I cells were transferred on day 0-2, and MRI was performed on day 6. For peak-stage transfer, OT-I cells were transferred on day 6 and mice were imaged on day 11. Adoptive transfer of OT-I cells had a therapeutic benefit in both early and peak stage paradigms (Fig 5A-D). For peak-stage treatment, in the turbinates (representative images shown in Fig 5A), the volume of bleeding decreased from 12.2 ± 1.3 mm^3^ to 1.6 ± 1.1 mm^3^ on day 11 (Fig 5E). In the OB (Fig 5B), the number of hypointensity spots decreased from 665 ± 127 to 114 ± 23 on day 11 (Fig 5F). In the brain (Fig 5D), the number of hypointensity spots decreased from 329 ± 72 to 83 ± 19 on day 11 (Fig 5G). T cell transfer reduced microbleeds (Fig 5B, C) and also reduced viral titers on day 6 in the OB. Relative VSV to actin RNA decreased from 0.95 ± 0.37 to 0.29 ± 0.07 (Fig 5H).

**Figure 5.**
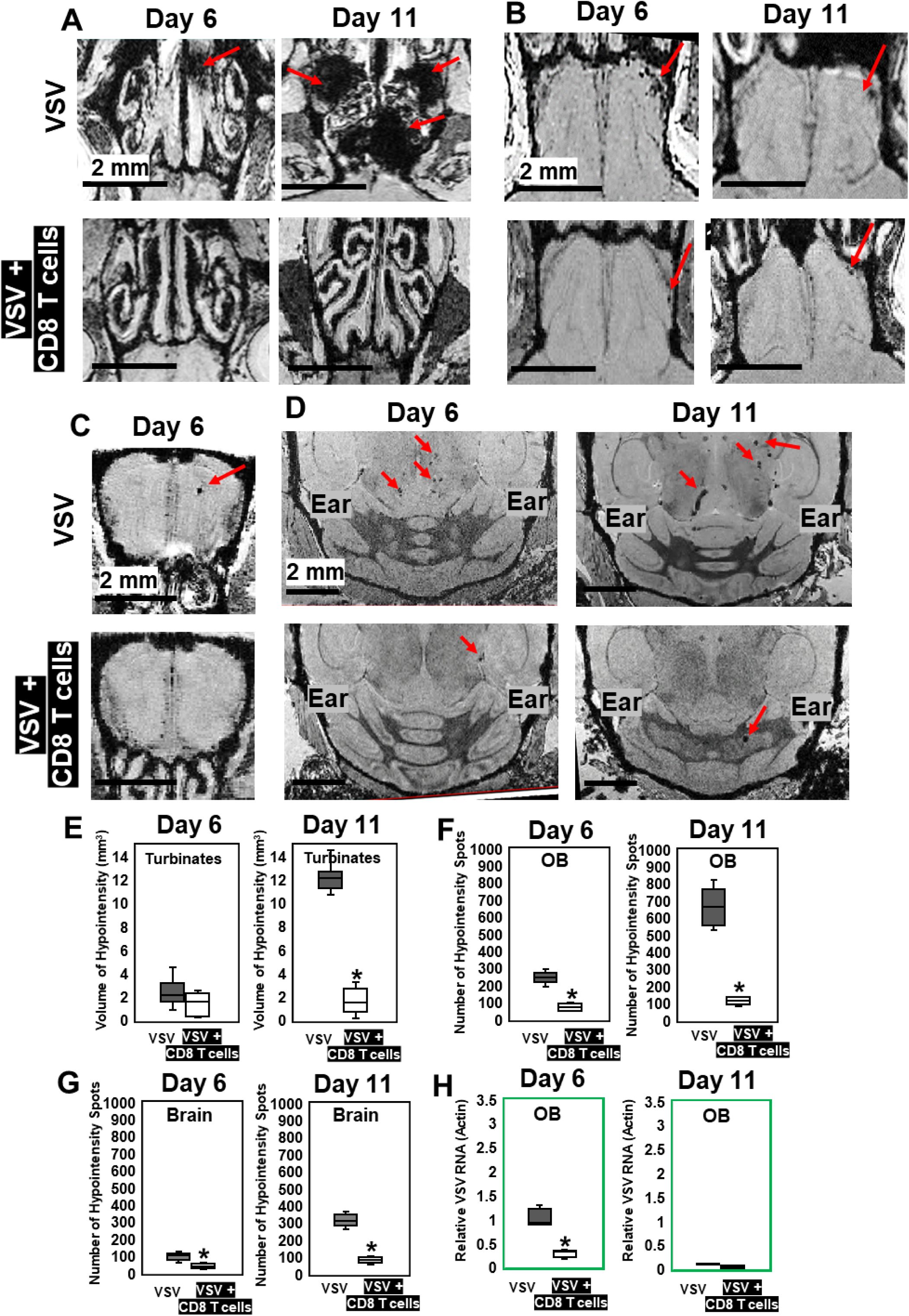
MRI study showed that adoptive CD8 T cells transfer reduced cerebrovasculature breakdown. (A-D) MRI showed that OT-1 CD8 T cells transfer reduced bleeds in the turbinates (A), GL of the OB (B), center of the OB (C) and brain (D) on day 6 and day 11. Red arrows, microbleeds. Quantification of volume of bleeding in the turbinates (E) and numbers of hypointensity spots in the OB (F) and brain (G) without or with T cells transfer on day 6 and day 11 (N = 5-7 mice per group). * p < 0.0001. (H) CD8 T cells reduced viral titers on day 6. *, p < 0.0001.

IHC confirmed that, without T cell transfer, there were a large amount of VSV and microbleeds, and a small number of CD8 T cells at the ONL, GL (Fig 6A) and center of the OB on day 6 (Fig 6D). Fig 6A and D also showed that GL and OB center were major sites of virus invasion and replication, which was consistent with Fig 1H. Upon transfer of OT-I cells, an increased number of CD8+ T cells, and a reduced amount of VSV and microbleeds was observed in the GL (Fig 6B) and OB center (Fig 6E). Quantification of these data is shown Fig 6C and F. Some microbleeds, however, were still seen in mice receiving antiviral T cells, especially in regions with a high density of T cells.

**Figure 6.**
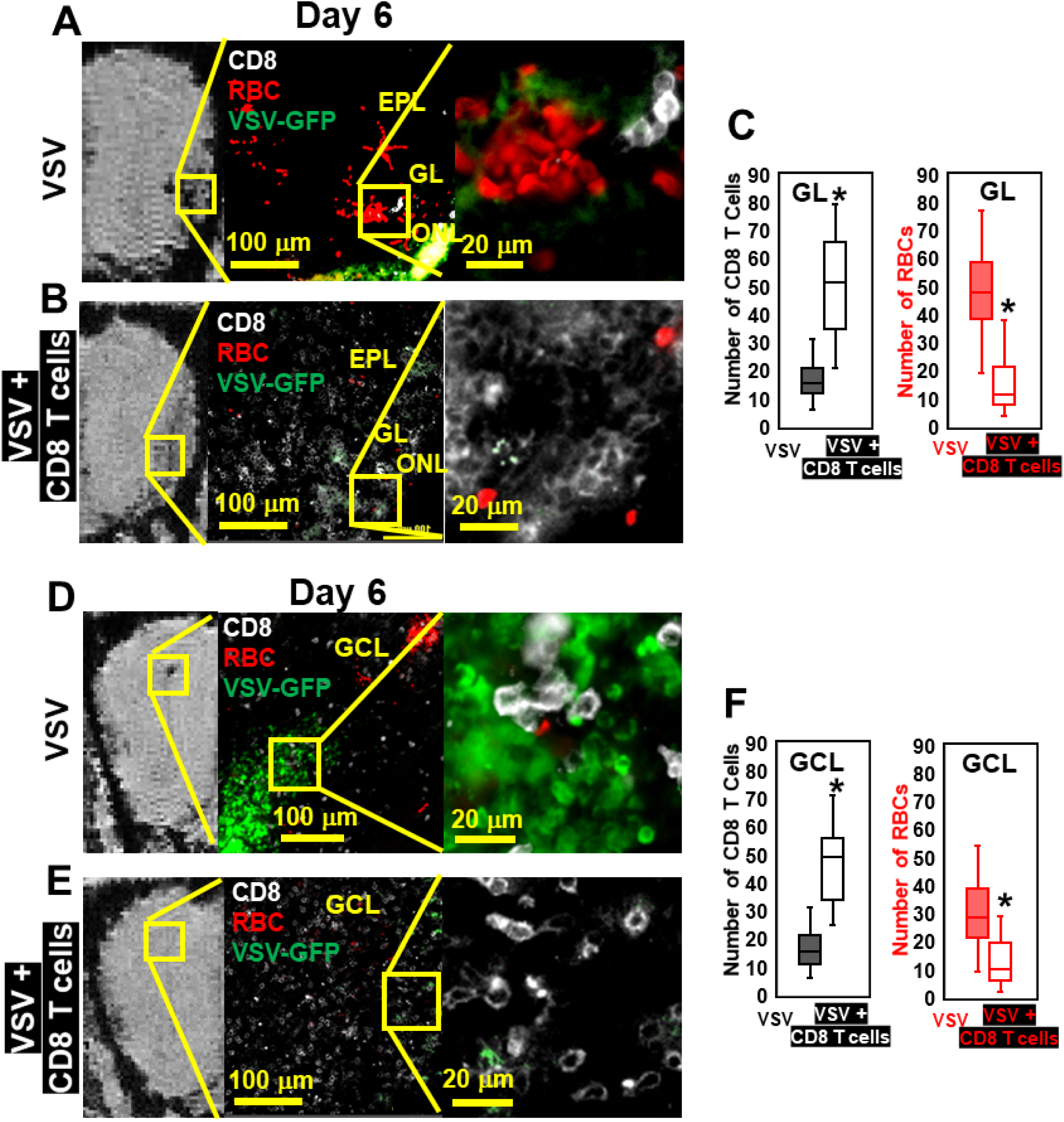
IHC study verified that antiviral CD8 T cells reduced VSV-induced brain bleeding. (A) On day 6, VSV induced bleeding at the ONL and GL and there was a small number of CD8 T cells. (B) CD8 T cells transfer reduced bleeding and virus and there was a larger number of CD8 T cells at the ONL and GL. (D, E) In the GCL, T cell transfer also reduced bleeding and virus comparing with non-T cells transferred. White, Brilliant Violet 421 conjugated anti-CD8 for T cells; Green, VSV-GFP; Red, Alexa Fluor 647 conjugated anti-Ter119 for RBCs. (C, F) Quantification of CD8 T cells and bleeds in the GL and GCL, under 40x view (0.2 mm x 0.2 mm). N = 4-6 mice per group. 6 views were counted manually per mouse. *, p < 0.0001.

### Individual MPIO labeled T cells can be detected by MRI

To understand the spatial relation between transferred T cells and microbleeds, we developed a method to label T cells with MPIO particles, which can be detected by MRI with high sensitivity (Pothayee et al., 2017; Shapiro et al., 2004; Wu et al., 2006). Labeling non-phagocytic T cells with contrast agents is challenging. Cationic agents, such as transfection agents and HIV-TAT peptide, or electroporation have been shown to increase the labeling efficiency (Bhatnagar et al., 2013; Hingorani et al., 2020; Thu et al., 2012). We therefore modified the surface of MPIO particles with cationic amine groups to increase the efficiency of uptake by T cells. A representative structure of the particle is shown in Fig 7A, and a detailed synthetic scheme is shown in Fig S6. Briefly, a streptavidin modified MPIO was used in this study. MPIO was chemically modified with -NH_2_ to increase uptake. In addition, anti-CD3 was conjugated to MPIO as an affinity ligand and to increase internalization of the particles by T cells. DyLight 488 was conjugated to MPIO for FACS and fluorescence microscopy studies.

**Figure 7.**
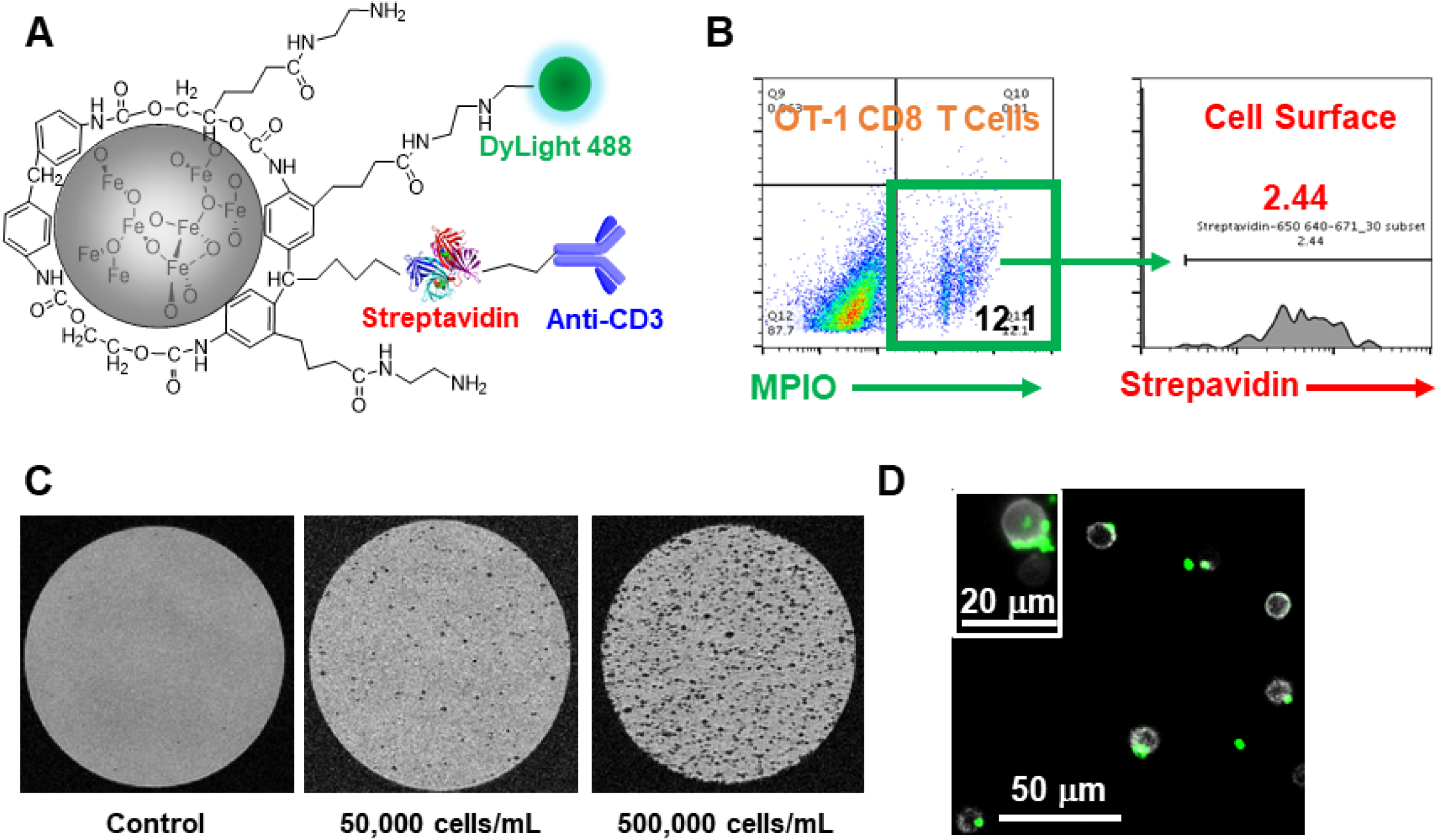
Label CD8 T cells with MPIO for MRI investigation. (A) A representative structure of the conjugated MPIO particle used in this study to label T cells. (B) FACS MPIO-labeled CD8 T cells, defined as DAPI^-^, mTomato^+^, DyLight 488^+^, DyLight 650^-^. (C) MR sensitivity of labeled T cells. (D) Fluorescence images of labeled T cells.

mTomato CD8+ OT-I T cells were incubated with MPIO overnight. For FACS analyses, anti-streptavidin was used to identify the location of MPIO as either extracellular (i.e., attached to the cell membrane) or intracellular. As shown in Figure 7B, T cell labeling efficiency was ∼12% and the majority of MPIOs were localized inside the cells. MPIO-labeled T cells showed MR sensitivity enabling single cell detection (Fig 7C) with some T cells containing more than one MPIO (Fig 7D). Labeling T cells with MPIO did not affect T cell function as measured by IL-2 and IFN-gamma release (Fig S7).

### MRI based tracking of T cells and microbleeds on day 1 of infection

With MRI having the sensitivity to detect both single T cells and microbleeds, we used this approach to guide histology studies and determine what gave rise to hypointensities at the earliest stages of VSV infection and inflammation in the brain. We wanted to determine whether vessel breakdown or T cell extravasation occurred first after adoptive transfer of antiviral T cells.

MPIO-labeled OT-I cells were transferred just prior to VSV infection. At 15-hr post-infection, no hypointensities were detected in the OB by MRI (Fig 8A). At 24-hr post-infection, the earliest hypointensity spots were detected near the GL (Fig 8B) and the center, MCL/GCL areas (Fig 8C). Similar hypointensities were not detected in the controls infused with unlabeled CD8 T cells. Since MPIOs and blood generate similar T2* effects, IHC was required to assess the cause of the hypointensities detected by MRI. IHC showed that, in the GL, CD8 T cell infiltration and bleeding co-localized and both occurred by 24 hr (Fig 8D). The MRI hypointensity in the GL was caused by the combination of MRI contrast from MPIO-labeled T cells and microbleeds. Interestingly, hypointensities in the MCL/GCL areas were due only to MPIO-labeled T cells, and no evidence for bleeding was detected (Fig 8E-G). To verify the presence of MPIO-Alexa Fluor 488 particles in transferred T cells, VSV with no GFP was used in the experiment shown in Fig 8D-F. The presence of VSV-GFP in the center areas on day 1 is shown in Fig 8G. As shown in Fig 1J and 4K, the center of the bulb was a major site of virus infection. It is likely that the ability of T cells to traffic to sites of viral infection rapidly prior to vascular damage explains why adoptive transfer of T cells greatly reduced brain bleeding during VSV infection. That T cells can migrate to sites of viral infection prior to vessel break down is consistent with earlier studies using the LCMV model to study this issue (Herz et al., 2015).

**Figure 8.**
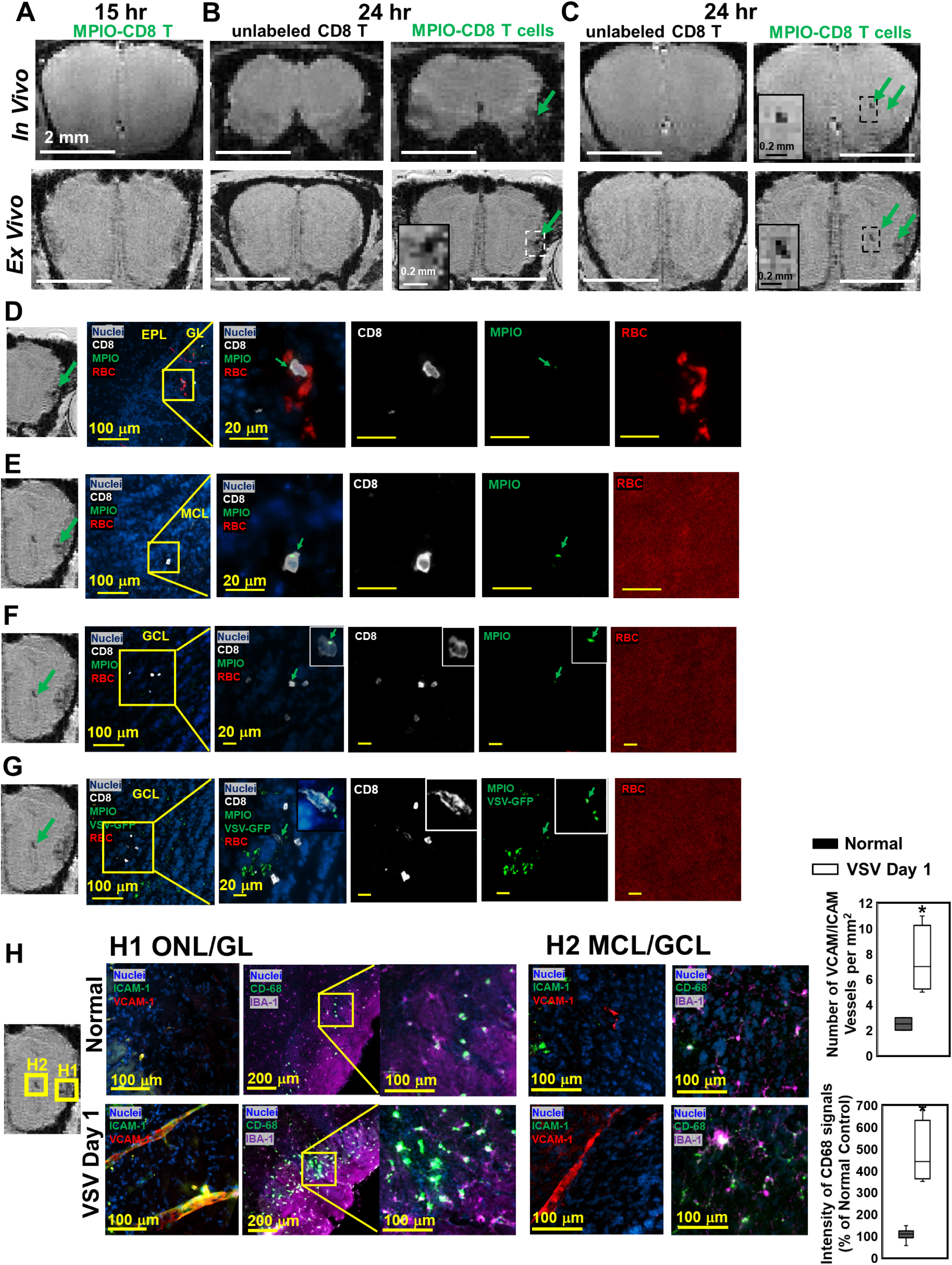
MRI tracking of virus-specific CD8 T cells and detection of brain viral infection on day 1. (A) No hypointensity was detected at 15-hr post-infection in the OB. (B, C) The earliest hypointensity spots were detected near the GL (B) and the center, MCL to GCL, (C) at 24-hr post-infection. Similar hypointensities were not detected in the controls infused with unlabeled CD8 T cells. RBCs were detected at the GL (D), but not near the MCL (E) and GCL (F, G), as shown by IHC. In (D-F), VSV with no GFP was used to verify the presence of MPIO-Alexa Fluor 488 in the T cells. White, Alexa Fluor 594 conjugated anti-CD8 for T cells. Green, VSV-GFP or MPIO-Alexa Fluor 488; Red, Alexa Fluor 700 conjugated anti-Ter119 for RBCs. (H) Activation of vessel endothelium cells and microglia were found at the ONL/GL (H1) and MCL/GCL (H2) on day 1. N = 3 per group. 3-6 views were quantified per mouse. *, p < 0.0001.

On day 1, at the locations that CD8 T cells and microbleeds were detected, activation of vascular endothelium cells (as indicated by VCAM-1 and ICAM-1) and microglia (as indicated by CD68) were also detected by IHC (Fig 8H). Thus, MRI of microbleeds and cell tracking provides a noninvasive method for early detection of cerebrovascular inflammation.

The combination of MRI contrast from MPIO-labeled T cells and bleeds increased MRI sensitivity to areas where there was both microbleeds and T cell infiltration (Fig 8B). To test whether this enabled better detection of sites of inflammation for later stages of infection, MRI hypointensity spots were counted after T cell transfer at day 6 and day 11 (Supplementary Results, Fig S8). More hypointensity spots could be detected likely due to additive effects of bleeding and T cell detection. The limitation at this stage is that we cannot distinguish microbleeds from MPIO-labeled T cells, and some of the MRI sites may be due to bleeding alone or T cell infiltration alone. However, the total signal from MPIO-labeled immune cells plus bleeds made sites of inflammation detectable by MRI. This provides an approach for imaging neuroinflammation when peripheral cells enter the brain, which has also been shown in previous studies focused on stroke and traumatic brain injury (Drieu et al., 2020; Foley et al., 2009).

## Discussion

Inflammatory processes are involved in most neurological disorders but are especially important during viral infections. Interactions between the immune system and CNS as well as the pathways by which immune cells infiltrate the brain differ depending on the anatomical location, structure of the barriers, and type of disorder. Using microbleeds as a marker of cerebrovasculature breakdown, we studied the relationship between vascular disruption and CD8 T cell infiltration using T2*-weighted MRI during a neurotropic VSV infection of the mouse brain.

T2*-weighted MRI (gradient-recalled echo or susceptibility-weighted imaging) exhibits a high sensitivity to microbleeds. This high sensitivity is due to paramagnetic properties of deoxy-hemoglobin from RBCs or its degradation product, hemosiderin (Haller et al., 2018). Using T2*-weighted MRI, paramagnetic and supermagnetic materials can create a magnetic field gradient that extends an MRI effect 50 times greater than the actual size of the material (Lauterbur, 1996). This amplification is critical for MRI detection of vascular effects during brain activation (BOLD functional MRI), for sensitive detection of bleeding by MRI, and for many pre-clinical studies that have labeled and imaged cells with MRI. Sensitivity down ∼1 pg iron per voxel can be achieved at high resolution. This is why we followed *in-vivo* MRI with *ex-vivo* MRI of fixed brains because the latter provides significantly higher resolution to guide histology. There is the potential to detect ∼10 deoxygenated RBCs, which facilitates high sensitivity visualization of vessel bleeds where only a small amount of blood has entered the brain. MRI was shown to be more sensitive than CT to detect bleeding. For example, it was demonstrated in a study of herpes simplex virus induced encephalitis that T2*-weighted MRI could detect bleeds in the brain within 2 days during the acute phases of the disease, whereas 1-2 weeks were required for CT-based detection (Gupta et al., 2012). The presence of microbleeds in patients with stroke and traumatic brain injury is increasingly recognized as an indicator of worse outcome (Griffin et al., 2019; Haller et al., 2018). Iron deposition can initiate continued neuroinflammation and brain damage. Indeed, MRI detected brain vessel damage and microbleeds in COVID patients was recently used to guide histological studies and uncover mechanisms of inflammation associated with SARS-CoV-2 (Lee et al., 2020; Poyiadji et al., 2020).

Guided by MRI we demonstrated that neurotropic VSV can cause brain vessel breakdown without the involvement of the peripheral immune system (Fig 3). VSV spike G-protein binds to low-density lipoprotein receptor (LDL-R) and enters cells via receptor-mediated endocytosis (Nikolic et al., 2018). It was reported that endothelium of brain capillaries express LDL-R (Meresse et al., 1989). In addition, a cell culture experiment showed that VSV-GFP infected and lysed endothelial cells (Patel et al., 2020). This may explain the presence of VSV along vessels and provides a potential mechanism for viral disruption of vascular integrity independent of peripheral immune cell infiltration into brain. Future studies are required to reveal the exact mechanism by which VSV infects endothelial cells and how this causes damage to cerebral blood vessels.

In our study we also demonstrated that the peripheral immune response to VSV infection is important for the maintenance of vascular integrity and pathogen clearance, as blockade of CNS immune cell entry via administration of anti-LFA1/VLA4 antibodies increased brain bleeding and viral titers (Fig 3). Moreover, supplementing the immune response by adoptively transferring activated virus-specific CD8+ T cells at either early (day 0-2) or later (day 6) time points post-infection protected cerebral vessels. Nevertheless, some bleeding was still observed even after virus-specific CD8+ T cells at days 0-2, and CD8+ T cells responding to the virus were associated with vessel damage at this early stage. Therefore, the transferred CD8+ T cells were not effective at completely blocking early viral entry from nasal olfactory sensory neurons into the OB.

VSV showed tropism for the neurons in the GL and GCL and transferred virus-specific CD8+ T cells first arrived in these two layers between 15 and 24 hrs post-infection. A previous study demonstrated that CD8+ T cells controlled neuronal virus in this model through engagements with cross-presenting microglia (Moseman et al., 2020). In our study, adoptive of virus-specific CD8+ T cells improved CNS viral control by reducing titers and preserving vascular integrity in the GL and GCL (Fig 5). It is also interesting that these two layers contain proliferating immature neurons in rodents (Batista-Brito et al., 2008; Lois & Alvarez-Buylla, 1994). These immature neurons are generated by precursor cells in the subventricular zone (SVZ) and traffic through the rostral migratory stream (RMS) to different layers of the OB, primarily the GL and GCL, where they differentiate into neurons (Batista-Brito et al., 2008; Lois & Alvarez-Buylla, 1994). Many viruses have a tropism for immature neurons (Li et al., 2016), and the presence of immature neurons in these two layers may in part explain the tropism of VSV.

We also observed that administering too many virus-specific CD8+ T cells (3 x 10^6^ T cells) on day 6 post-infection can be pathogenic (Fig S9). This higher dose of cells (relative to the lower 5 x 10^5^ dose) led to increased cerebral bleeding. MRI showed that in the nasal turbinates both doses of T cells reduced bleeding. However, in the OB, midbrain, and hindbrain, the higher dose of CD8+ T cells caused significantly more bleeding compared with the lower dose. This finding was confirmed via immunohistochemistry and suggests that antiviral CD8+ T cells can contribute to vessel damage while attempting to control virus. CD8+ T cell mediated vascular pathology was also observed during cerebral malaria (Swanson et al., 2016). Thus, care must be used when determining T cell doses for therapeutic adoptive cell transfers.

In conclusion, our MRI guided studies have revealed that both viral infection and CD8+ T cells can contribute to cerebrovascular bleeding following a nasal viral infection. VSV likely has the ability through direct infection of cerebrovascular endothelium to cause vascular damage; however, our high dose T cell transfer studies show that having too many antiviral T cells can also elevate cerebral bleeding. While additional studies are required to identify specific viral and immune mechanisms that result in vascular disruption, it is clear that control of a CNS viral infection requires maintenance of a precise balance between reducing titers and preserving tissue. Enhancement of adaptive immunity can come at the cost of increased cerebrovascular pathology. MRI can aid in our assessment of CNS viral infections and the efficacy of the therapies we use to intervene. Development of more sensitive MRI contrast agents that can be distinguished from blood will be crucial for future immune cell tracking studies and should facilitate our ability to detect neuroinflammation (Barbic et al., 2021; Zabow et al., 2014). Lastly, additional studies of pathogen-specific T cells via MRI should advance our understanding T cell entry and movement throughout the brain, which will aid in optimizing adoptive cell therapies for the treatment of infections and tumors (e.g., anti-cancer chimeric antigen receptor T cell therapy).

## Materials and Methods

### Animal procedures

C57BL/6J (B6), B6.129(Cg)-Gt(ROSA)26Sortm4(ACTB-tdTomato,-EGFP)Luo/J (mTomato), and C57BL/6-Tg(TcraTcrb)1100Mjb/J (OT-I) mice were purchased from The Jackson Laboratory. mTomato x OT-I mice were bred and maintained under specific pathogen-free conditions at the National Institutes of Health. All mice in this study were handled in accordance with the Institute of Laboratory Research guidelines and were approved by the Animal Care and Use Committee of the National Institute of Neurological Disorders and Stroke.

### Intranasal (i.n.) VSV Infection

Twelve-to 13-week-old C57BL/6 mice were used in all the infection and control experiments. Recombinant VSV expressed a truncated form of OVA and with or without GFP. For intranasal (i.n.) VSV-OVA infection, a dose of 3.5 x 10^4^ PFU in 10 μL of PBS was pipetted into the opening of each nostril.

### MRI study

For MRI scans, VSV infected mice were placed into the chamber shown in Figure S1. MRI experiments were performed at 11.7 T on a Bruker Biospec MRI system (Bruker BioSpin). T2*-weighted 3D gradient-recalled echo (GRE) sequences were used for acquisitions. For *in-vivo* imaging, the following parameters were used: isotropic resolution = 75 μm, TE/TR = 10/30 ms, FA = 10°, NA = 3. After *in-vivo* acquisitions, mice were perfused transcardially with 5% formalin. The heads were post-fixed with 10% formalin. 24-hr before *ex-vivo* MRI imaging acquisition, heads were transferred to PBS buffer. For *ex-vivo* MRI: isotropic resolution = 50 μm, TE/TR = 20/40 ms, FA = 15°, NA = 12.

### Quantification of bleeding in the nasal turbinates and brain

*Ex-vivo* high-resolution MRI was used to quantify nasal and brain bleeding in VSV infected mice. Because bleeding in the turbinates was so extensive, volumes were measured. Hypointensity volumes were obtained by manually drawn serial voxels of interest (VOI) using the Medical Image Processing, Analysis, and Visualization (MIPAV) program (http://mipav.cit.nih.gov) (Saar et al., 2015). Hypointensities in the brain were more focal so the number of spots was counted manually. The volume of hypointensities and number of hypointensity spots from normal healthy mice were used as background and subtracted from the measurements for each infected mouse.

### Quantitative real-time PCR (RT-qPCR) for VSV titers

Total RNA was isolated from OB using miRNeasy Mini Kit (Qiagen). Subsequently, 1 μg of RNA was treated with amplification grade DNAse I (Life Technologies) to remove any contaminating genomic DNA. We used 250 ng of total RNA for cDNA synthesis using iScript cDNA Synthesis Kit (Bio-Rad). All Q-PCRs were performed in 20 μL volumes using SYBR Green PCR master mix (Applied Biosystems) in a 96-well optic tray using a CFX96 Real-Time PCR machine (Bio-Rad). The reactions were conducted in duplicate, and samples without reverse transcriptase were used as non-template controls. Forward primer (specific for the VSV genes): TGA TGA TGC ATG ATC CAG CTC T; reverse primer: ACA CAC CTC CAA TGG AAG GGT. Actin RNA was used as internal standard. Q-PCR reactions were run with an initial denaturation temperature of 95 °C for 10 min, followed by 40 cycles of three-step amplification (denaturation at 95 °C for 10 s, 60 °C annealing for 15s and extension at 72 °C for 20 s).

### Adhesion molecule blockade studies

Anti-LFA-1 (clone M17/4), anti-VLA-4 (clone PS/2), and isotype control (rat IgG2a) antibodies were purchased from BioXcell. Mice were treated with a combination of 500 μg anti-LFA-1 and 500 μg anti-VLA-4, or 1000 μg isotype control antibody at day 1, 3, and 5 post-infection through intraperitoneal injection. Normal healthy mice were used as the control group.

### Adoptive transfer of CD8 T cells for MRI

We considered day 6 post-infection as the peak of encephalitis in this model of VSV infection (Huneycutt et al., 1994). For MRI studies ranging from day 1 to day 6 post-infection, we injected 5 x 10^5^ mTomato OT-I CD8+ T cells into the tail vein on day 0 or 2. T cells were injected intravenously on day 6 for MRI studies performed on day 7 to 11. Figure S1 provides an illustration of these two experiment paradigms.

### OT-I CD8 T cell isolation, culture, and labeling with MPIO

Splenocytes were isolated from mTomato OT-I mice as described in the Supplemental Methods. OT-I CD8 T cells were isolated using a Dynabeads Untouched Mouse CD8 Cells Kit (Invitrogen). The purity of CD8 T cells was confirmed by flow cytometry (Fig S5), using the Beckman Coulter’s MoFlo Astrios cell sorter and analyzed by the Summit 6.3.1 software. CD8 T cells were cultured overnight in RPMI medium supplemented with OVA_257-264_ peptide (2 μg/ml), 2-mercaptoethanol (50 μM), FBS (10%), penicillin (50 U/ml), streptomycin (50 μg/ml), and l-glutamine (2 mM) and incubated at 37 °C with 5% CO_2_ in a humidified (85% humidity) cell culture incubator.

For MPIO labeling, mTomato OT-I CD8+ T cells were incubated with MPIO at a ratio of 1:5 overnight. The CD8 T cells were then washed with cell staining buffer (BioLegend), harvested, and stained with anti-streptavidin-DyLight 650 and DAPI for 30 min on ice. Anti-streptavidin-DyLight 650 was used to identify the cellular location of MPIO, since MPIO was modified with streptavidin. MPIO labeled OT-I cells, defined as DAPI^-^, mTomato^+^, DyLight 488^+^, DyLight 650^-^, were collected. We used the Beckman Coulter’s MoFlo Astrios cell sorter installed with the Summit 6.3.1 software for cell sorting.

### Bioconjugation of MPIO

We generated MPIO-NH_2_-anti-CD3-DyLight 488 particles to label and track mTomato OT-I CD8+ T cells by MRI. The synthetic method and scheme are shown in the Supplementary Methods (Fig S6).

### Histology

Following perfusion or *ex-vivo* MRI acquisition, brains were isolated and post-fixed with 5% formalin for 2 hr. Brains were then transferred into 15% sucrose for 1 day, followed by 30% sucrose for 1 day. Brains were cut coronally and embedded in tissue freezing medium (Fisher HealthCare). Cryosections were cut coronally (30 μm) by a cryostat (Leica CM 1850). Brain sections were stained using a standard procedure for free-floating immunohistochemistry. Briefly, brain sections were rinsed with TBS 3 x 5 min, followed by blocking in 5% normal serum in TBS with 0.5% Triton X-100 for 2 hours at 4 °C. Then the sections were stained in the following antibody solutions in TBS with 3% normal serum and 0.3% Triton X-100 overnight at 4 °C. To label red blood cells (RBC), Alexa Fluor 647 or Alexa Fluor 700 conjugated anti-TER119 (2.5 μg/mL; BioLegend) was used. For vascular endothelial cells, Alexa Fluor 594 or Alexa Fluor 647 conjugated anti-mouse CD31 (2 μg/mL; clone MEC13.3; BioLegend) was used. To study vessel inflammation, Alexa Fluor 488 anti-CD54 antibody (2 μg/mL, ICAM-1, YN1/1.7.4, BioLegend), and Alexa Fluor 647 anti-CD106 antibody (2 μg/mL, VCAM-1, 429 MVCAM.A, BioLegend) were used. For CD8 T cells, Brilliant Violet 421 or Alexa Fluor 594 conjugated anti-mouse CD8a (2 μg/mL; clone 53-6.7; BioLegend) was used. To study microglia and activation, rabbit anti-IBA-1 antibody (0.4 μg/mL, FUJIFILM Wako) and rat anti-CD68 antibody (5 μg/mL, clone FA-11, Bio-Rad) was used. For secondary antibody staining, Alexa Fluor 700 goat anti-rabbit or Alexa Fluor 488 goat anti-rat were used, and the staining was at 4 °C for 2 hrs. Then tissue sections were put on the Superfrost microscope slides (Fisherbrand). After dried overnight at room temperature, slides with brain sections were then coverslipped using ProLong Gold antifade reagent with or without DAPI (Thermo Fisher Scientific). Images were captured using a Nikon Eclipse Ti microscope (Nikon).

### Quantification of vessel VCAM-1 and ICAM-1 coverage and intensity of CD68 signals

Images obtained under 20x view (0.4 mm x 0.4 mm) were used for quantification. ImageJ was used for quantification. During quantifications of vessel coverage, the CD31 area was taken as 100% and the VCAM-1 or ICAM-1 positive area was expressed as a percentage normalized to CD31 area.

### Statistical analysis

Statistical analysis was carried out with the Student’s t-test. A p < 0.05 was considered statistically significant.

## Acknowledgements

This research was supported by the intramural program at the National Institute of Neurological Disorders and Stroke (NINDS), National Institutes of Health (NIH). The authors would like to thank Dr. Jasmin Herz for the valuable discussion during the development of T cell labeling and tracking method.

## Author Contributions

L.L. performed the experiments and wrote the manuscript with input from other authors. S.D. performed MRI experiments. R.H. and N.P. prepared samples for *ex-vivo* MRI and contributed to *ex-vivo* quantification of microbleeds. T.A. contributed to the T cell labeling method. N.B. performed T cells transfer. D.M. performed flow cytometry analysis and cell sorting. E.A.M. and S.G. contributed to sample preparation for *ex-vivo* MRI, intranasal infection, and viral titer experiments. D.B.M. designed experiments and wrote manuscript. A.P.K. supervised the whole project and wrote manuscript.

## Supplementary Methods

### Splenocytes isolation

Splenocytes were prepared by smashing with the textured end of a sterile 10-mL syringe followed by pipetting through a 40-micron filter into a 15-mL conical tube. The cells were collected by centrifugation at 300 g for 6 min and decanted. Cell pellets were resuspended in mouse ACK lysis buffer and waited for 10 min at room temperature followed by two washes with PBS (w/1% FBS).

### MPIO bioconjugation for T cell labelling

MPIO particles with streptavidin modification and carboxylic acid group on the surface (Dynabeads™ MyOne™ Streptavidin C1, Thermo Fisher Scientific) was used for the bioconjugation reactions. Conjugation reactions were performed as shown in Supplementary Figure 4. 0.5 ml of MPIO particles (5 x 10^9^) was resuspended in 10 ml of 2-(N-morpholino)ethanesulfonic acid buffer (MES, 50 mM, pH 5.5). 1-Ethyl-3-[3-dimethylaminopropyl]carbodiimide hydrochloride (EDAC) and sulfo-N-hydroxysuccinimide (Sulfo-NHS) were added to the particle suspension. The final concentrations of EDAC and NHS were 75 mM and 150 mM, respectively. The reaction mixture was stirred at room temperature for 30 min. The resulting NHS ester was purified by a magnetic stand separator (Miltenyi Biotec). 20 μl of ethylenediamine (shall be large folds excess of calculated carboxyl groups on the particles) was dissolved in 10 ml PBS and pH was adjusted to 7.5. The purified NHS ester was added to the ethylenediamine solution. The reaction mixture was allowed to stir for 4 hr at room temperature. The resulted MPIO-NH_2_ particles were purified by a magnetic separator and washed 2 times with borate buffer (50 mM, pH 8.5, with 0.05% Tween 20). 50 μg of DyLight 488 or DyLight 650-NHS ester (Thermo Fisher Scientific) was added to MPIO-NH_2_ particles in 5 ml of borate buffer. Covered the vial with aluminum foil to prevent photobleaching and placed it on a slowly rotating platform for 1 hr at room temperature. Wash the beads 3 times with antibody washing and binding buffer (AWBB, 50 mM Tris-HCl, 150 mM NaCl, 0.05% Tween 20, pH 8.0). 25 μl Biotin anti-mouse CD3 stock solution (12.5 μg, BioLegend) was added to 5 ml of the beads in AWBB. Covered the tube with aluminum foil and placed it on the rotating platform for 30 min at room temperature. The resulted particles were washed and kept in AWBB. Right before T cell labeling, beads were washed 3 times with RPMI medium.

### IL-2 and IFN-gamma release assay for T cell function

Briefly, MPIO-labeled OT-1 CD8 T cells or unlabeled CD8 T cells (1 x 10^6^) were treated with brefeldin A (BioLegend) for 4 hrs. Then cells were stained with APC anti-mouse IFN-γ and PE/Cy7 anti-mouse IL-2 antibodies (BioLegend), by using the Cyto-Fast Fix/Perm Buffer Set (BioLegend) followed the Intracellular Flow Cytometry Staining Protocol from BioLegend.

## Supplementary Results

### The combination of MRI contrast from MPIO-labeled T cells and bleeds increased sensitivity to microbleeds

Upon transfer with unlabeled CD8 T cells, on day 6 post-infection, MRI could detect localized hypointensities (*in-vivo*) and punctate hypointensities (*ex-vivo*) at the edge of OB, near GL and EPL (red arrows in Fig S8A). These hypointensities were caused by bleedings. Upon transfer with MPIO-labeled CD8 T cells, more hypointensities were detected, not only near the edge, but also in the center of the OB. These hypointensities were caused by microbleeds, plus the infiltration of MPIO-labeled T cells (red and green arrows in Fig S8B). As quantified in Fig S8C and D, significantly more hypointensity spots from the mice transferred with MPIO-labeled T cells were detected than the mice transferred with unlabeled T cells in the OB and brain. The number of hypointensty spots in the OB transferred with unlabeled T cell and MPIO-labeled T cell were 72 ± 22 and 124 ± 20 on day 6, respectively (Fig S8C). When CD8 T cells were transferred on day 6, on day 11, the difference in number and intensity of the hypointensity spots between the two groups was even more significant. The combination of MRI contrast from MPIO-labeled T cells and bleeds increased sensitivity to microbleeds about 2 folds at each time point (Fig S8C, D). Representative IHC image verified the infiltration of MPIO-labeled OT-1 CD8 T cells in the brain. Due to the similarity in contrast at this stage we cannot distinguish microbleeds from MPIO-labeled T cells. However, the total signal from MPIO-labeled T cells plus bleeds made sites of inflammation easily detectable by MRI, which might provide an approach for imaging neuroinflammation.

---

**Supplementary Figure 1.**
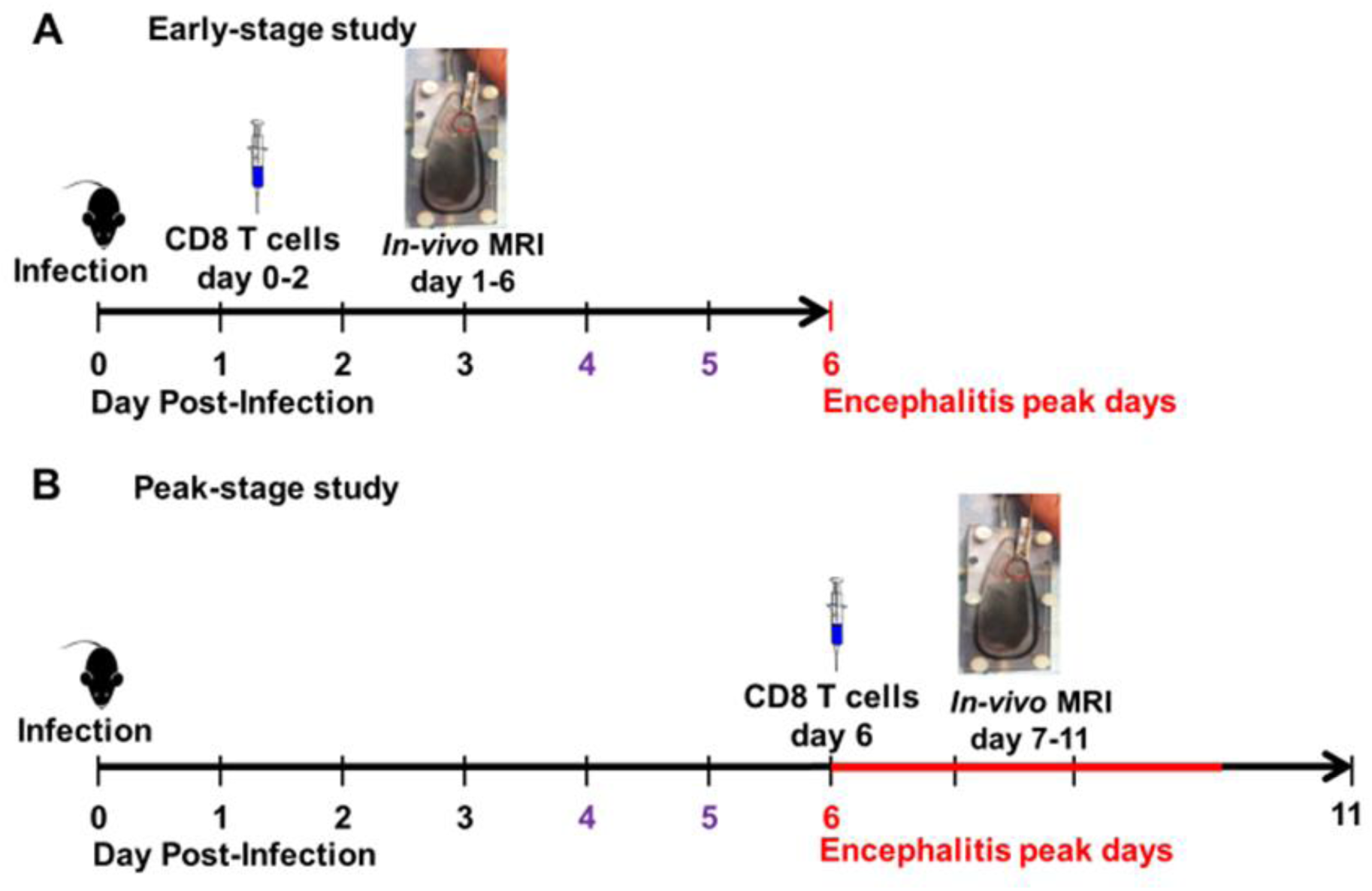
Experiment paradigms. (A) Paradigm for CD8 T cell transfer study during early-stage infection. CD8 T cells were transferred on day 0-2. *In-vivo* MRI study was performed from day 1 to 6. The encephalitis peak time was day 6, which was labeled in red. Remarkable amount of CD45+ cells infiltration into the OB was reported since day 4, which was labeled in purple. (B) Paradigm for CD8 T cell transfer study during peak-stage infection. CD8 T cells were transferred on day 6. *In-vivo* MRI study was performed from day 7 to 11.

**Supplementary Figure 2.**
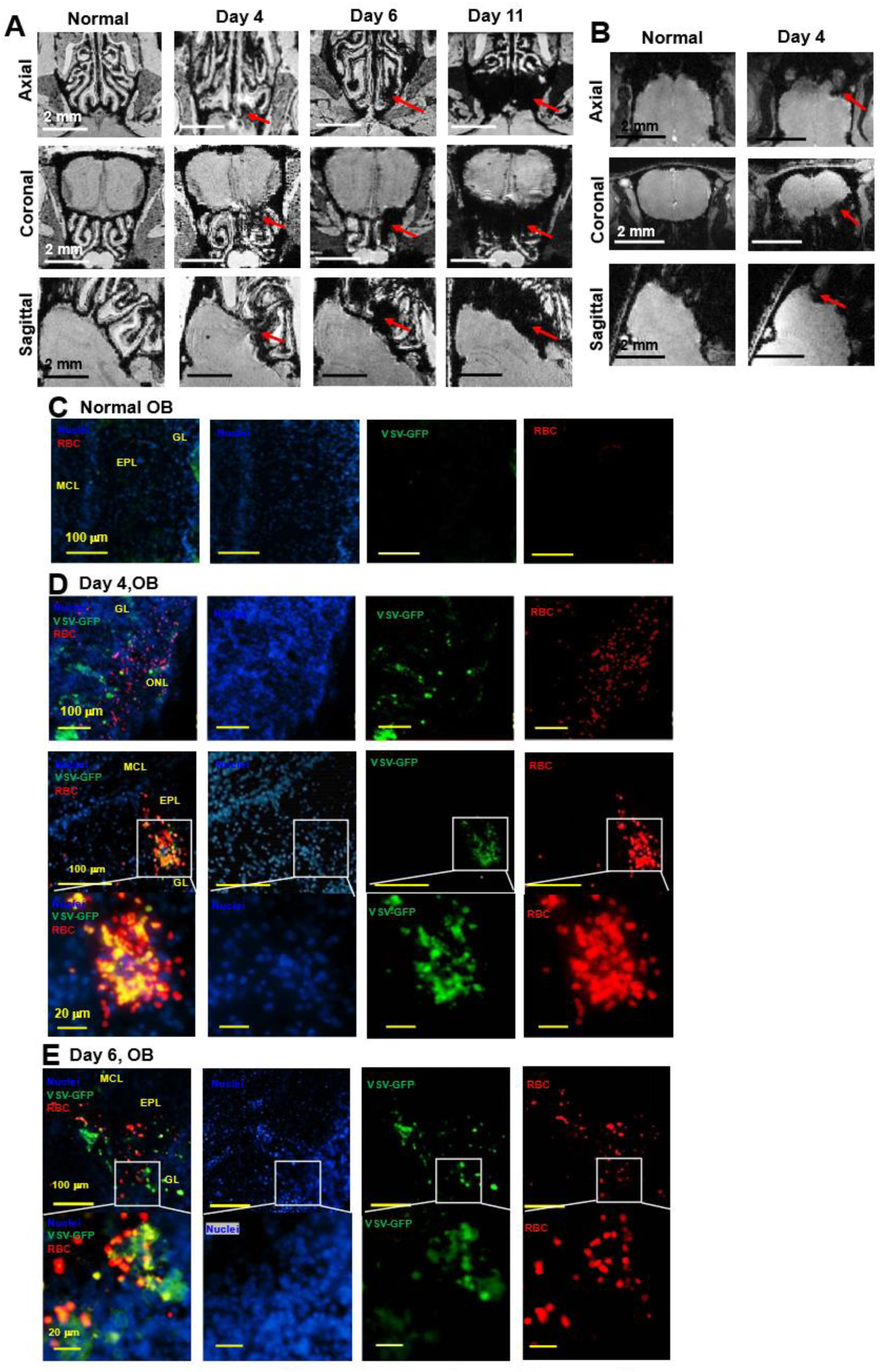
MRI detected microbleeds in the turbinate and OB during VSV infection. (A) 3D view of *ex-vivo* MRM of turbinates from normal mouse and VSV-infected mice on day 4, 6, and 11. (B) 3D view of *in-vivo* MR images of OB from normal mouse and VSV-infected mice on day 4. (C-E) IHC verification of the MRI results. Separated color channels of fluorescence images of the OB from a normal mouse (C) and VSV-infected mice on day 4 (D) and 6 (E). GL, glomerular layer; EPL, external plexiform layer; MCL, mitral cell layer. Blue, DAPI for nuclei; Green, VSV-GFP; Red, Alexa Fluor 647-conjugated anti-Ter119 for RBCs.

**Supplementary Figure 3.**
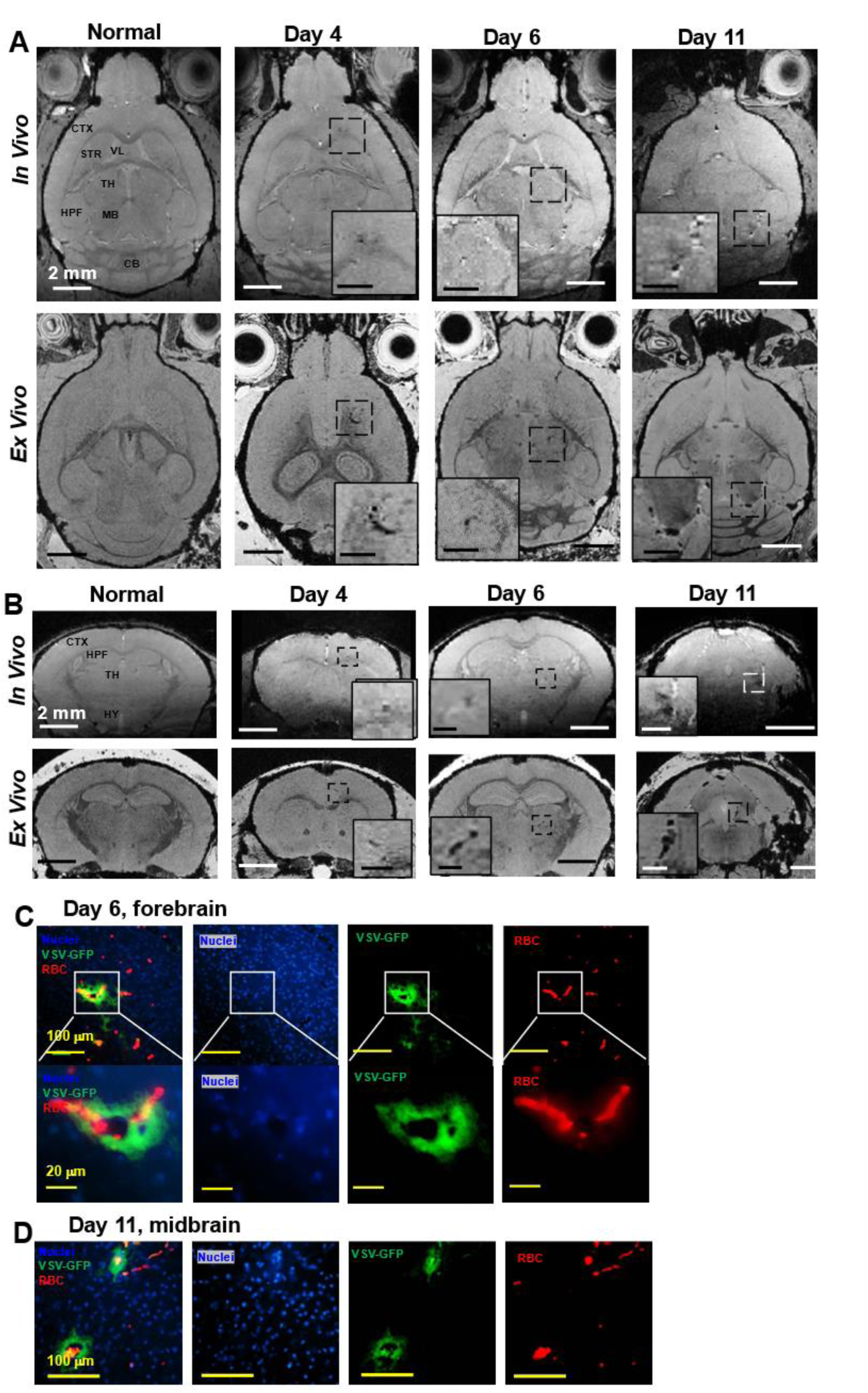
MRI monitored vessel breakdown in the rest of brain. More (A) axial view and (B) coronal view of *in-vivo* (upper panel) and *ex-vivo* (lower panel) MR images of brain, showing the breakdown of vessels from frontal brain to mid-brain. (C, D) IHC verification of the MRI results. Separated color channels of fluorescence images to show microbleeds and VSV in the forebrain on day 6 (C) and midbrain on day 11 (D) . Blue, DAPI for nuclei; Green, VSV-GFP; Red, Alexa Fluor 647-conjugated anti-Ter119 for RBCs.

**Supplementary Figure 4.**
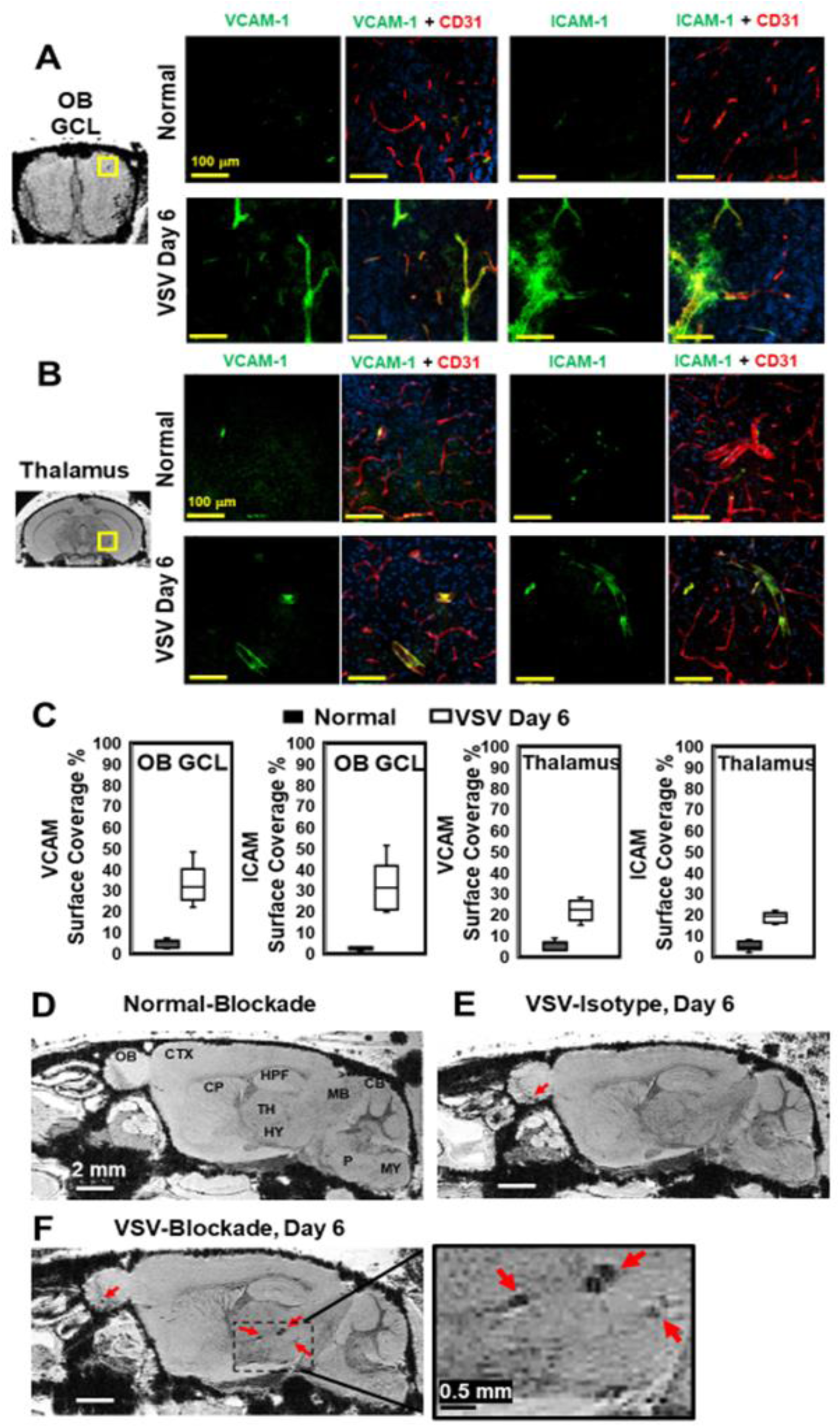
Sagittal view of MR images of the brain showed a large area of hemorrhage at the thalamus of anti-LFA-1/VLA-4 treated mouse on day 6.

**Supplementary Figure 5.**
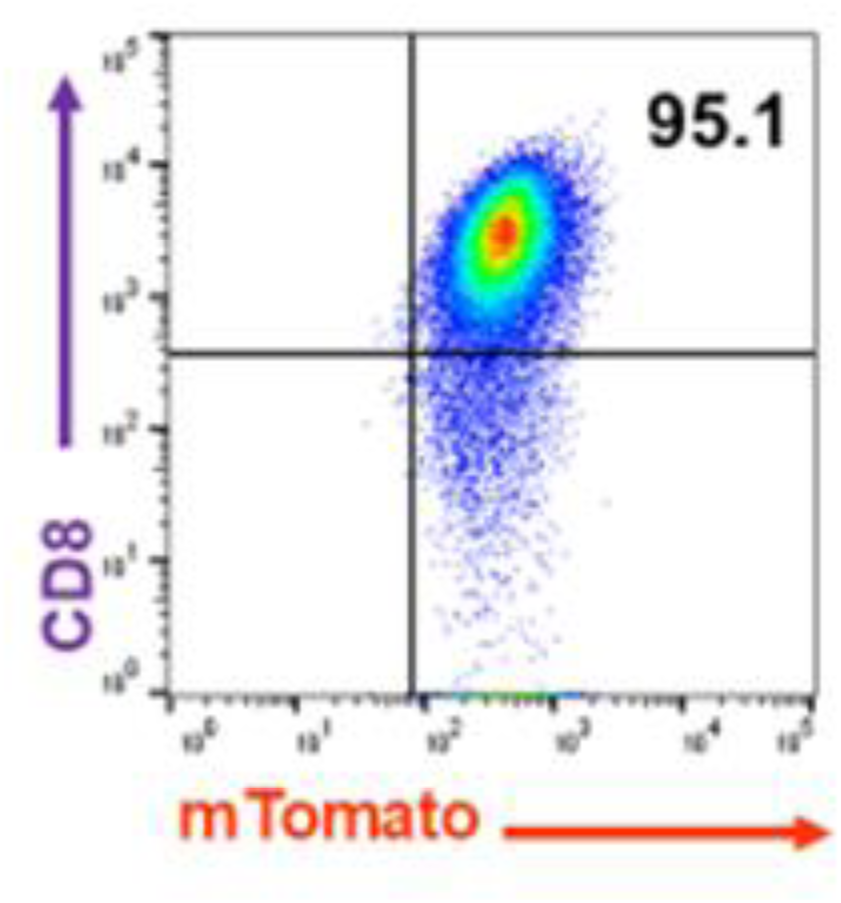
Purity of CD8 T cells used in this study. CD8 T cells were isolated using the Dynabeads Untouched Mouse CD8 Cells Kit.

**Supplementary Figure 6.**
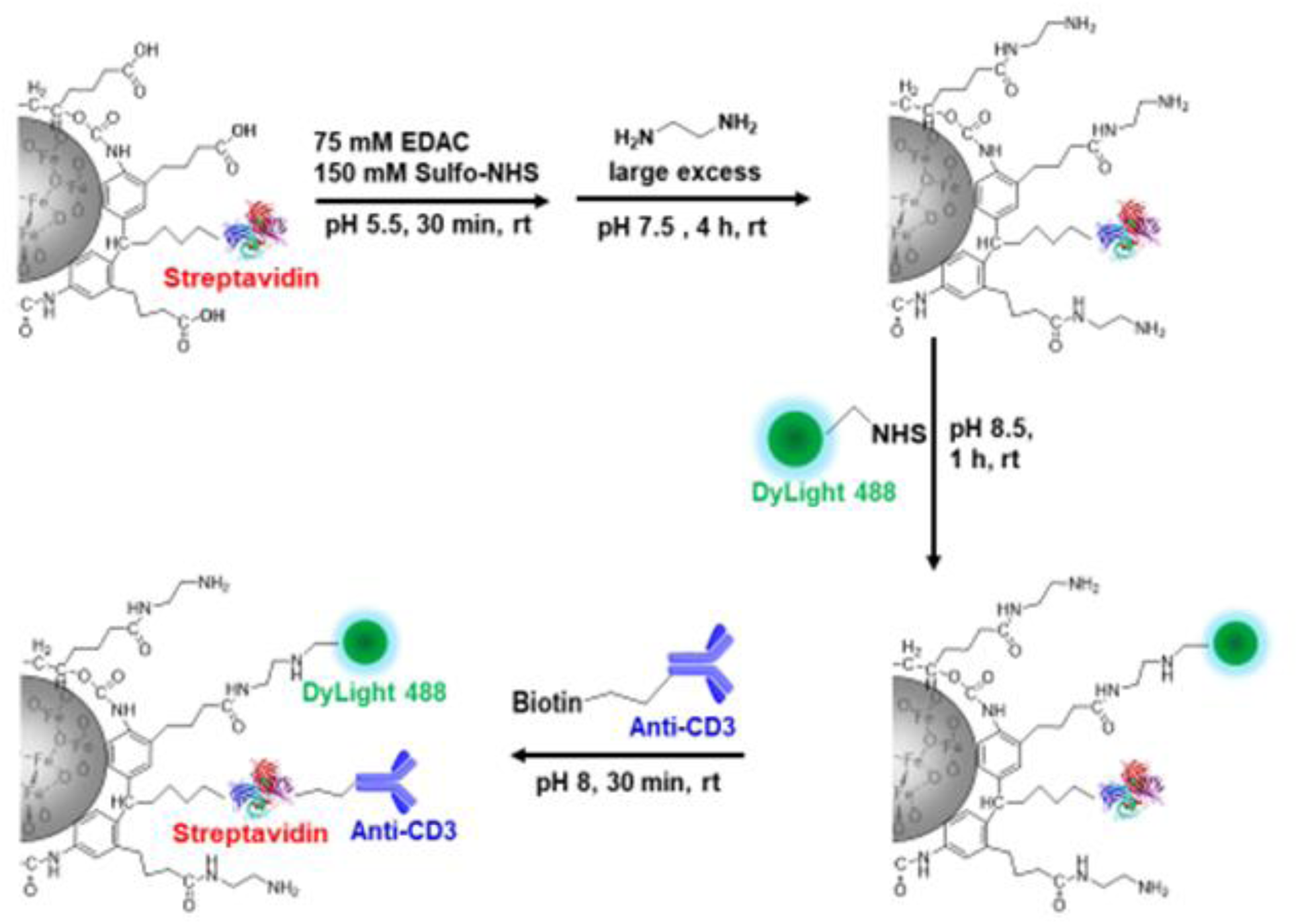
Bioconjugation of MPIO particles to label T cells.

**Supplementary Figure 7.**
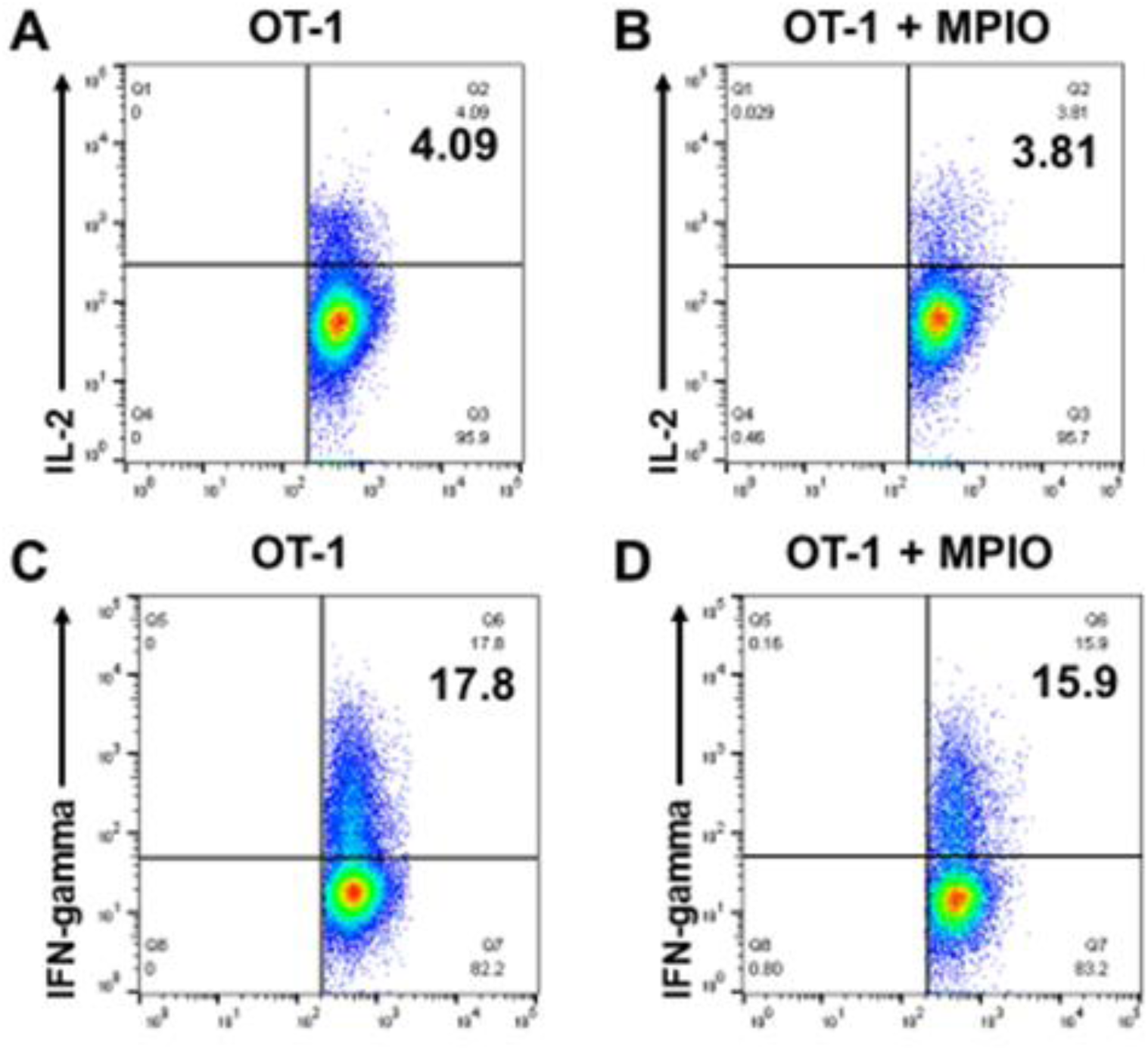
Labeling T cells with MPIO did not affect T cell function on IL-2 and IFN-gamma release.

**Supplementary Figure 8.**
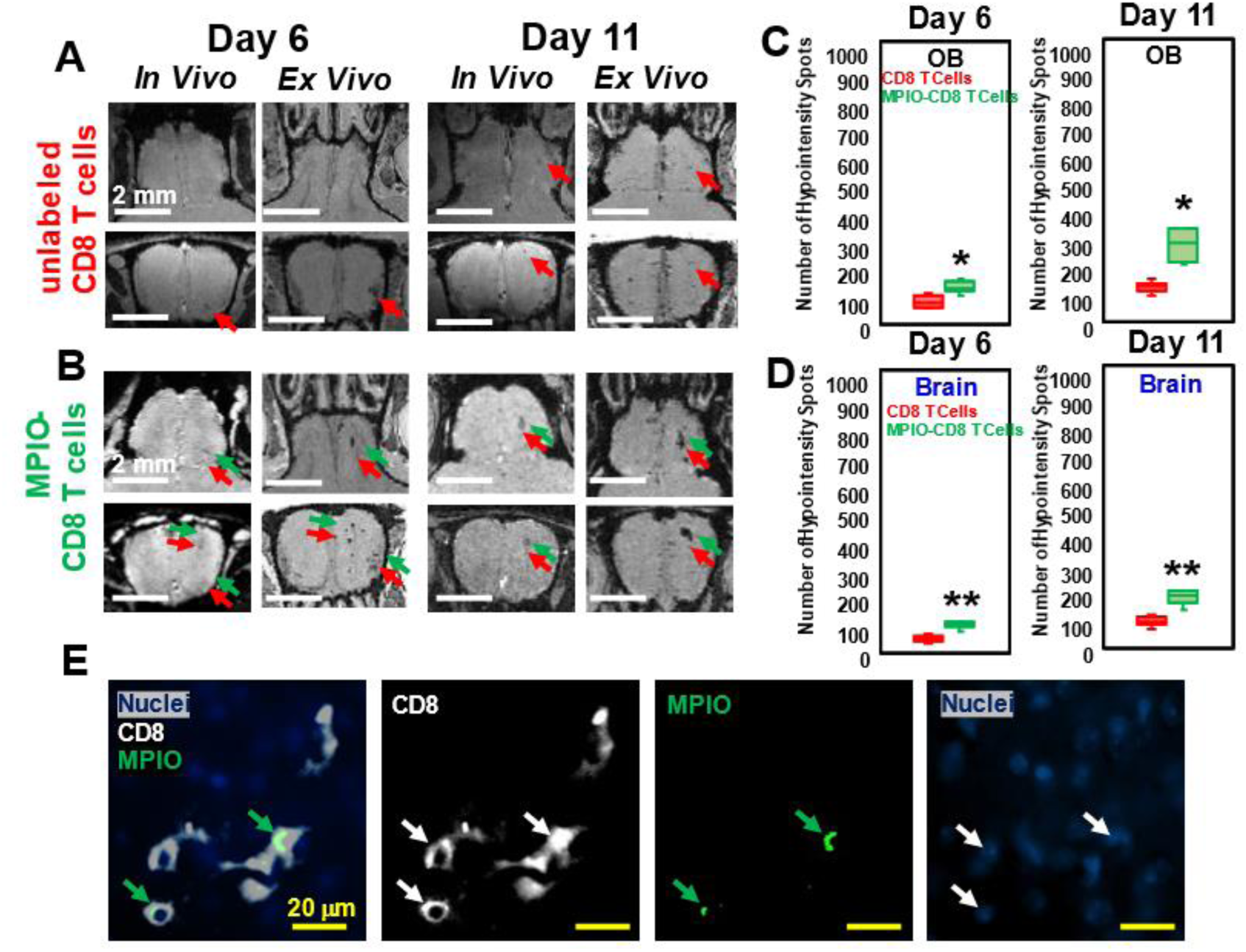
MPIO-labeled T cells increased MRI sensitivity to microbleeds. (A) MRI, on day 6 and 11, of OB from the mice transferred with unlabeled CD8 T cells. (B) MRI from the mice transferred with MPIO-labeled CD8 T cells. Red arrows, hypointensities caused by microbleeds. Green arrows, hypointensities caused by infiltration of MPIO-labeled CD8 T cells. Quantification of the number of hypointensity spots in OB (C) and brain (D) on day 6 and 11, upon transfer of unlabeled CD8 T cells (N = 5 mice per group) or MPIO-labeled CD8 T cells (N = 7 mice per group). Red bar, quantification from transfer of unlabeled T cells. Green bar, quantification from transfer of MPIO-labeled T cells. * p < 0.001; ** p < 0.0001. (E) IHC study showed the infiltration of MPIO-labeled CD8 T cells. Green arrows, MPIO. Blue, nuclei; White, CD8 T cell; Green, MPIO.

**Supplementary Figure 9.**
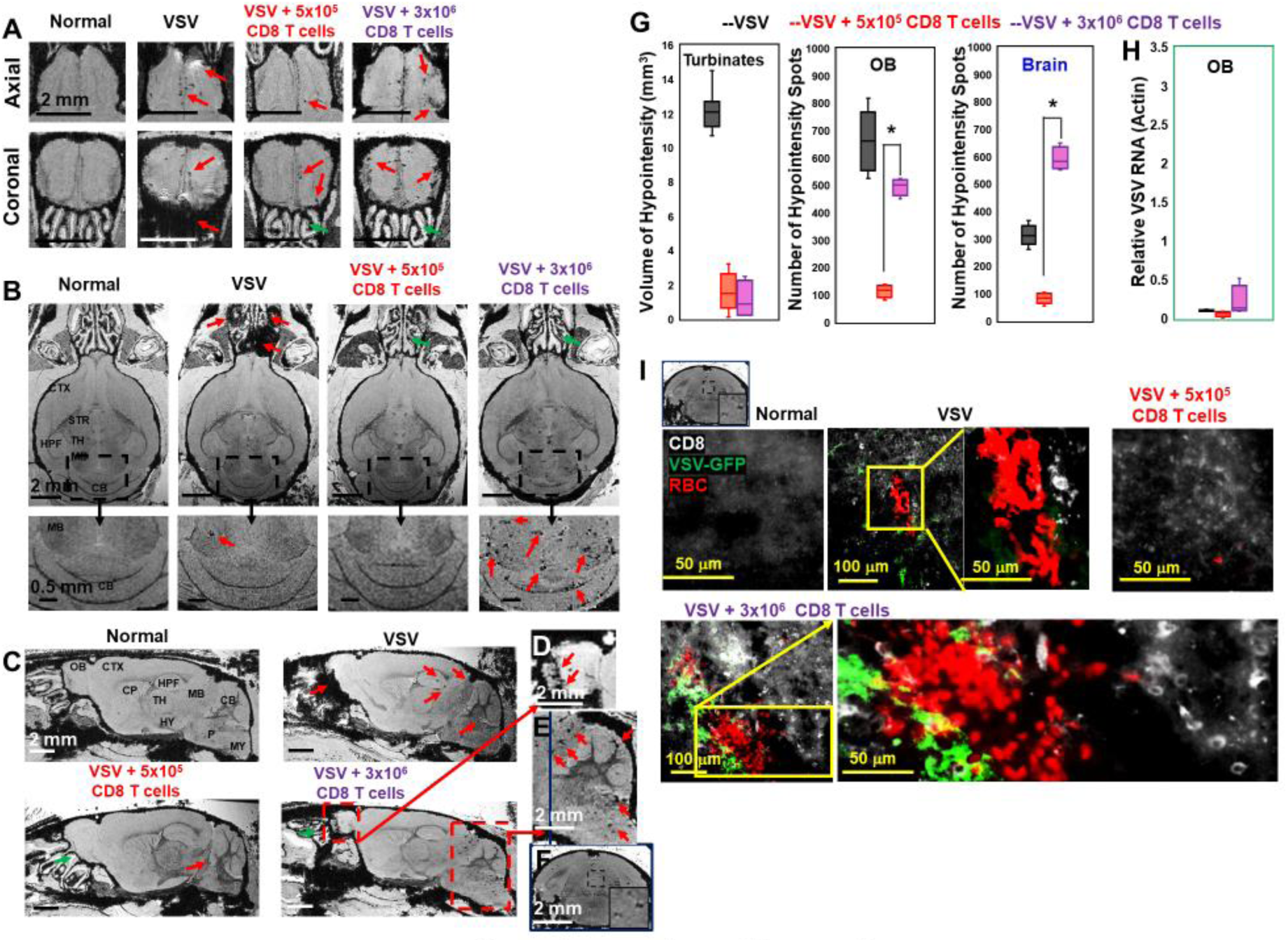
Adoptive transfer of overdose of CD8 T cells (3 x 10^6^ T cells) at the peak stage caused bleeding in the OB and brain. T cells are transferred on day 6 and images were shown for day 11. Axial views (A), coronal views (B), and sagittal views (C) of normal mice, VSV infected mice with no T cells transfer, with the transfer of 5 x 10^5^ CD8 T cells, with the transfer of 3 x 10^6^ CD8 T cells. (D, E, F) were enlarged views. Green arrows, reduced bleeding in the turbinates. Red arrows, bleeding in the turbinates, OB, or brain. (G) Quantification of bleeding volume in turbinates, and number of hypointensity spots in OB and brain without (black bar, N = 7) or with T cell transfer (red bar, 5 x 10^5^ T cells, N = 5; purple bar, 3 x 10^6^ T cells, N = 5) on day 11. * p < 0.0001 comparing with 5 x 10^5^ T cells. (H) Viral titer. (I) IHC study showed large amount of VSV, CD8 T cells, and RBCs in the midbrain of overdose treated mice. Green, VSV-GFP; White, OT-1 CD8 T cells; Red, Ter119 for RBCs.

